# Disentangling the contribution of trait plasticity to improve the productivity of a maize-soybean intercrop system for the Midwest, USA

**DOI:** 10.1101/2025.03.14.643314

**Authors:** Elena A. Pelech, Jochem B. Evers, Carl J. Bernacchi

**Author notes:** Corresponding Authors: Jochem Evers and Carl Bernacchi. Funding Information: Global Change and Photosynthesis Research Unit of the USDA Agricultural Research Service. Author contributions: Pelech collected plant data, parametrized and modified the existing model developed by Evers, ran simulations and wrote the first manuscript draft. Evers and Bernacchi supervised experimental design and writing.

## Abstract

Crop yields in intercropping systems are the result of a combination of factors dominated by plastic responses in plant traits to the heterogeneity associated with the intercrop design and the row configuration of the intercrop. Disentangling their relative influence is infeasible *in situ* but crucial for cultivar selection and intercrop design. Using functional-structural plant (FSP) modelling, these effects can be separated in silico. Here, a mechanistic FSP model was developed, including three-dimensional aboveground plant architecture of maize and soybean, radiation distribution, and assimilate allocation. The model was used to explore the potential to improve yields in a simultaneous intercrop by disentangling the contribution of three plastic traits related to photosynthesis, leaf thickness and plant height. The improved phenotypes were then simulated in two intercrop configurations, single- and twin-rows of maize, for potential increases in land-use efficiency. The study revealed that for maize, photosynthesis had the greatest contribution (+78%), followed by plant height (+31%) and leaf thickness (+6%) where the total maize monoculture phenotype produced the greatest maize yield without affecting the yield of intercropped soybean. However, soybean trait plasticity had a negligible effect on soybean yield, but the soybean monoculture phenotype with a low light-saturated photosynthetic rate resulted in the greatest intercropped maize yield. These improved phenotypes may increase land-use efficiency by 1-3% relative to the standard monoculture systems of the Midwest, USA. Together, these results could aid the selection of maize and soybean germplasm that could improve the productivity of a simultaneous intercrop.

## Introduction

Plant phenotypic plasticity, the capacity of a genotype to express different phenotypes in various environments (Sultan, 2000), drives numerous ecologically important traits related to physiology and performance to cope with heterogenous environments (Schneider, 2022; Sultan, 2000; Valladares et al., 2007). It is commonly assumed that phenotypic plasticity is advantageous for intercropping systems in realising yield advantages over monoculture systems of each component crop (Elhakeem et al., 2019; Postma & Lynch, 2012; Tilman & Snell-Rood, 2014). Studies have demonstrated that some plastic responses can be neutral to crop yield in simultaneous intercrop systems (Li et al., 2020), but beneficial in relay intercrop systems where different species are planted at different times which shortens the co-growth period (Zhu et al., 2015). Therefore, the assumption that phenotypic plasticity is advantageous depends on whether individual plastic traits overcome maladaptive competition within a specific intercrop system (Bongers et al., 2025).

Analysing the effect of individual trait plasticity on plant performance is difficult *in vivo*. Continuous feedback loops exist between plants and their local environment. The changed plant trait can also affect the environmental factors that drive plasticity, and multiple plastic traits can interact (Evers et al., 2019). Therefore, environmental heterogeneity is greater in intercrops compared to monocultures. Intercropping N-fixing legumes between wider than typical maize rows while maintaining typical maize planting density has captured the attention of USA farmers (Deichman and Kremer, 2019). The intercrop design, defined as the solar corridor, may improve yields by maximising the availability of incident light deeper into the canopy (Deichman, 2000; Deichman and Kremer, 2019; Kremer and Deichman, 2016). A previous study showed that maize exhibited physiological plasticity in response to growth in a solar corridor while soybean exhibited both physiological and architectural plasticity (Pelech et al., 2023). However, these responses were detrimental in light- and land-use efficiency for the solar corridor intercrop compared to the standard monoculture systems of the Midwest, USA. Disentangling the contribution of individual plastic traits to yield in this study and how these contributions depend on intercrop row configuration would highlight the most influential traits to intercrop performance.

Resolving the relative contributions of various plasticity responses would require inducing each plastic trait response independently while preventing plasticity in other traits in real plants, which is implausible to achieve *in situ* at this present time. Creating three-dimensional (3-D) virtual plants overcomes this challenge, allowing for investigation of variation in single and combined trait plasticity. The value of this approach has already been demonstrated by simulating crop systems to understand the contributions of individual plastic traits to plant performance (Evers et al., 2019; Gaudio et al., 2019). Functional-structural plant (FSP) models can be parameterised at the organ level to simulate plant growth in response to environmental variations while considering plant architecture in 3-D and physiological processes such as photosynthesis and assimilate allocation (Evers et al., 2018; Louarn & Song, 2020; Vos et al., 2010). Thus, the modelling framework can create virtual phenotypes where individual traits can be varied one by one, and the effect of each plastic trait on plant yield is quantified separately and in combination across multiple row configurations. Descriptive FSP models have been used to quantify the contribution of trait plasticity to light interception in intercrop systems but they did not consider the underlying physiological consequences (Barillot et al., 2014; Li et al., 2021; Zhu et al., 2015). Plant structure in these studies was an input, not an emergent property of the model based on the underlying physiological mechanisms governing plant growth and development. A mechanistic model with a representation of photosynthesis and assimilate distribution within each plant structure according to light interception and organ demands is a necessary step forward to quantify the contribution of plastic traits to yield.

This study parametrised a mechanistic maize and soybean FSP model with a representation of photosynthesis and assimilate distribution using data from a field experiment (Pelech et al., 2023). The model was then used to explore the potential for yield improvement in a simultaneous solar corridor intercrop simulation by disentangling the contribution three plastic traits. Plasticity in this study was defined as the difference in trait values for photosynthesis, leaf thickness and plant height; found to be significantly different between intercropped and monoculture systems for both maize and soybean (Pelech et al., 2023). The first objective quantified the role of plasticity to maize yield in both monoculture and intercrop configurations while keeping the soybean phenotype constant in the intercrop as maize is the main cash crop. For the second objective, individual plastic soybean traits were evaluated in the intercrop only once the highest-yielding maize phenotype was determined. And the third objective evaluated the improved maize and soybean phenotypes in the single-row and twin-row intercrop configurations by using the land equivalent ratio (LER), which is a measure for land-use efficiency of mixed crop systems. This research will ultimately allow selection of maize and soybean germplasm that could improve productivity in the solar corridor intercrop system compared to the standard monoculture systems of the Midwest, USA.

## Methods

### Model development

In the 2019 field experiment in Pelech et al., 2023 monoculture and intercrop plant data were collected on (i) individual leaf appearance and senescence at the phytomer level, (ii) plant organ dimensions and biomass at the phytomer level, and (iii) photosynthetic light response curves on the youngest fully expanded leaves. Air and soil temperature measurements were also collected throughout the 2019 field experiment for the calculation of thermal time (°C d, degree days) and daily modelled temperatures (*T_ave_*),

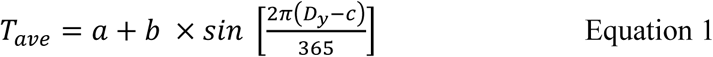

where *T_ave_* is the daily mean temperature (°C), *D_y_* is the day of year, *a* is the lowest temperature of the year (°C), *b* is the scale, and *c* is the day of year when temperature was the highest. The temperature probes (model 107, Campbell Scientific, Logan, UT, USA) recorded measurements every 15 s and were averaged over 30 minutes using a data logger (model CR1000 Campbell Scientific, Logan, UT, USA). Two temperature probes, one 0.35 m above the soil surface within the canopy and one in the soil at a 0.05 m depth, were placed between two rows in one plot of each crop system. Measurements began one week after germination, where temperature data from a nearby weather station was used to fill the measurement gap between planting and germination. Phenological development was then modelled assuming a base temperature of 10°C for both species (https://mrcc.illinois.edu/CLIMATE (Supplementary Table 1 and Supplementary Figures 1-2)

The FSP model was constructed based on an existing model (Evers & Bastiaans, 2016) using the GroIMP software package (Hemmerling et al., 2008). The model time step is equal to one day and includes three main parts 1) a dynamic representation of the 3-D architecture of maize and soybean plants, 2) a radiation model to simulate light capture of individual organs, and 3) a photosynthesis and assimilate distribution model. Simulations began with emergence and ended immediately before the onset of senescence. Therefore, whole plant senescence was not simulated and yield at the end of simulation represented yield at full physiological maturity. Canopy snapshots and comparison of the maize-soybean solar corridor intercrop from the 2019 field experiment versus a simulated scene at the same growth stage are shown in Figures 1-2.

**Figure 1.**
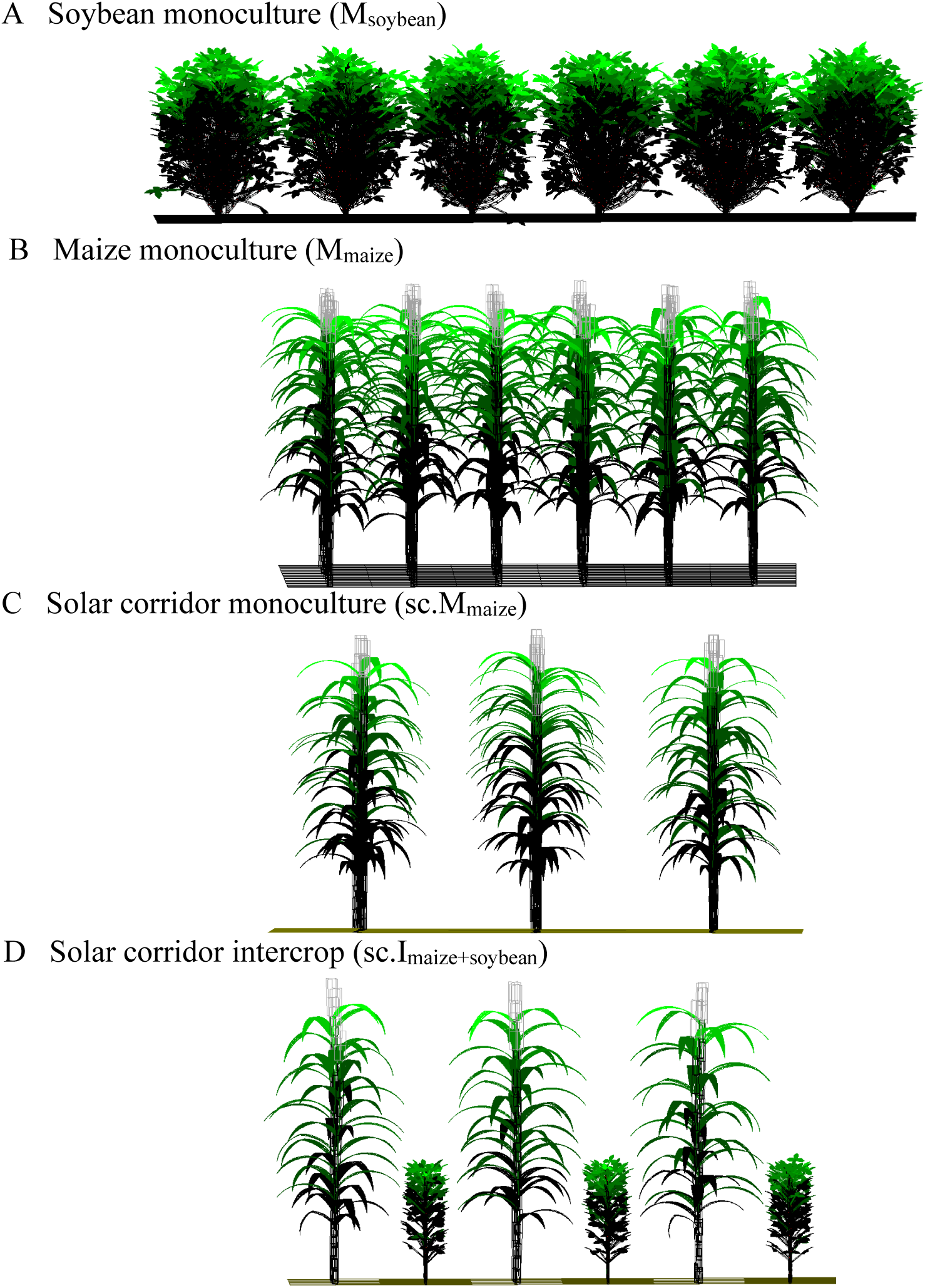
Model canopy snapshots of A soybean monoculture (M_soybean_), B maize monoculture (M_maize_), solar corridor monoculture (sc.M_maize_), solar corridor intercrop (sc.I_maize+soybean_). Each snapshot is at day 90, where the model time step is equal to one day. The colour gradients represent the proportion of light intercepted, where light interception increases from black to green for plant organs and from black to yellow for the soil surface. The figure is not to scale.

**Figure 2.**
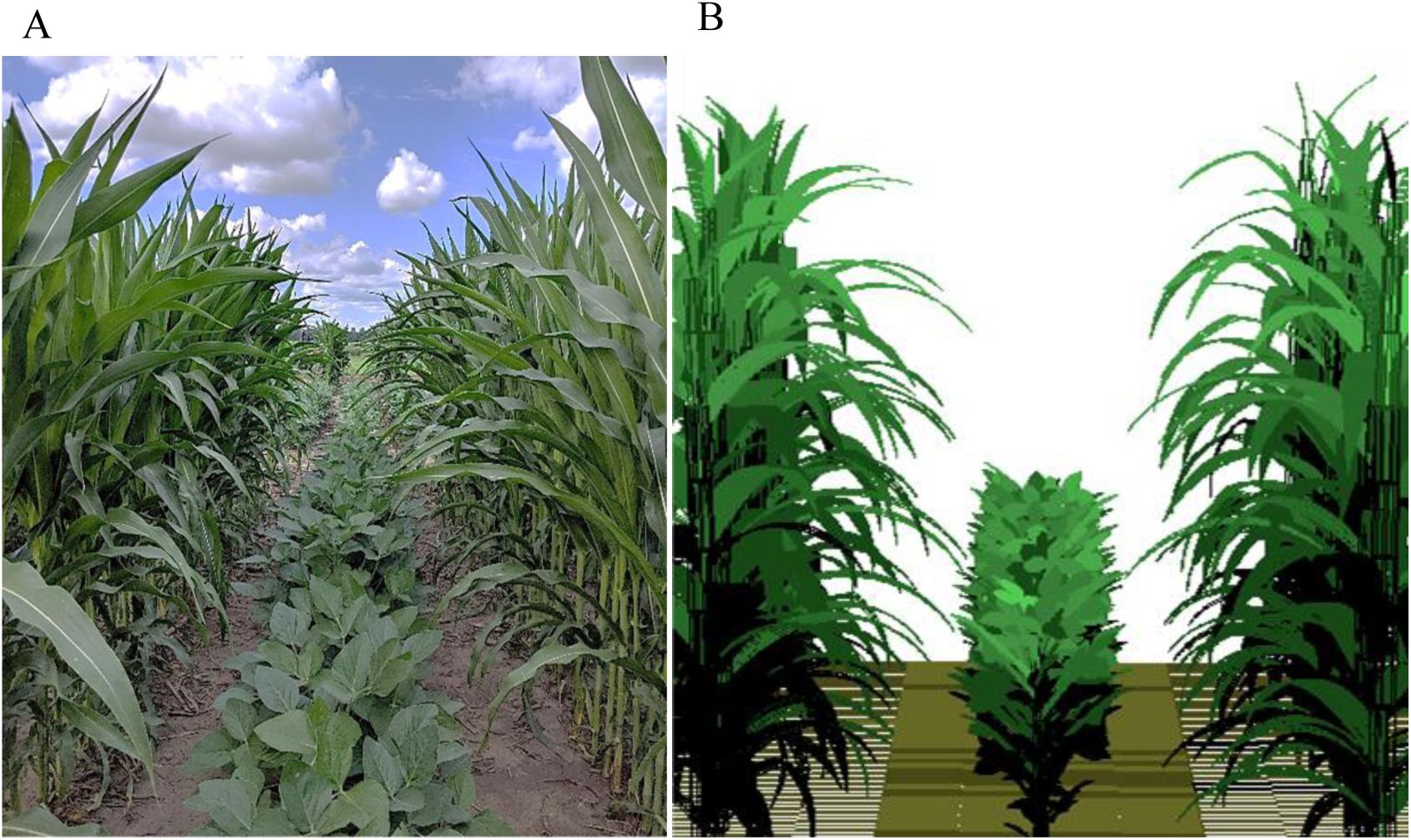
Comparison of A, maize-soybean solar corridor intercrop plot with B, a simulated scene at the same growth stage (42 days after emergence). The colour gradients in B represent the proportion of light intercepted, where light interception increases from black to green for plant organs and from black to yellow for the soil surface. The seasonal dynamics in field view are shown in Supplementary Video 4 and snapshots, Supplementary Figure 10.

**Figure 3.**
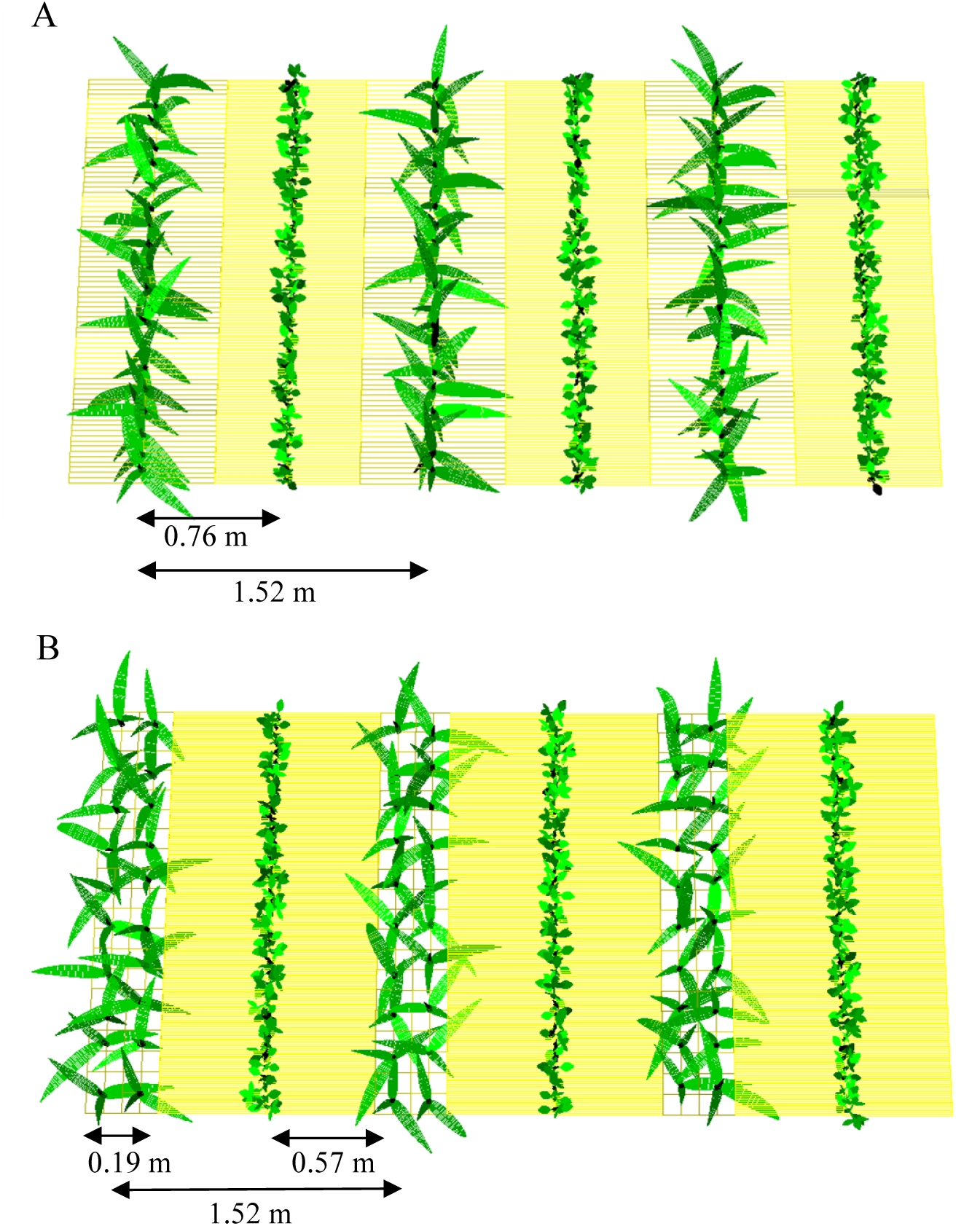
The comparison of A, single-row and B, twin-row arrangement of maize in the solar corridor intercrop plots at 15 days after emergence. The arrows represent the row dimensions. Both simulation plots had a cloned area of 100 m^2^. The colour gradients represent the proportion of light intercepted, where light interception increases from black to green for plant organs and from black to yellow for the soil surface. The figure is not to scale.

#### 1. The 3-D maize and soybean model

The model simulates 3-D plant architecture at the phytomer level, an architectural ranking unit consisting of individual plant organs in development which includes internode, sheath, leaf blade, and lateral ear for maize, and internode, axillary pods, petiole, petiolules and a trifoliate leaf for soybean. The model also considered two significant features of organ development: (1) the rates and duration of organ initiation, appearance, and elongation, and (2) geometric attributes. All plant input parameters are described in Supplementary Table 1. Both maize and soybean leaf geometric attributes were parameterised manually, except for leaf length-to-width ratio (Supplementary Figure 3) and soybean leaf insertion angle was calculated by image analysis in ImageJ (https://imagej.net/software/imagej/ [Supplementary Figure 4]). The senescence of individual leaves at the lower phytomers were simulated by inputting the average leaf life value calculated as the thermal time between a fully developed phytomer and leaf senescence (Supplementary Figure 2).

The first three simulated phytomers in maize had internode extension turned off, and the blade phyllotaxis represented a whorl. The remaining maize phytomers had internode extension turned on and represented distichous phyllotaxis i.e., the formation of successive leaves at 180° to each other. The maize model simulated determinate growth where maximum seed biomass and growth duration were initiated after 17 vegetative phytomers were reached. For soybean, the first phytomer represented the unifoliate leaves, and the remaining upper phytomer ranks represented trifoliate leaves. The phyllotaxis of the upper soybean phytomers was parameterised manually and the curvature of petioles was not considered. Soybean branching was enforced for monoculture simulations once individual plants reached eight phytomers, where the number of branches depended on the plant’s source/sink ratio. As the number of soybean branches was low in the intercrop (Supplementary Figure 4), branching was turned off to represent the intercrop phenotype of soybean. The soybean model simulates indeterminate growth where the time of flowering (48 days after emergence [DAE]) was a model input to ‘switch on’ the source/sink demand of flower (pod) organs while new phytomers continued to develop.

#### 2. The radiation model

The GroIMP radiation model was used to simulate light distribution and absorption by leaf organs. Both direct and diffuse radiation was simulated as described previously (Evers et al., 2010; Zhu et al., 2015). The direct incoming radiation was simulated using 24 equally dispersed directional light sources to represent the course of the sun. The position of each light source was modified by the day length, azimuth and solar elevation angle using Illinois latitude (40.6°) and starting day of year ([DOY] 163; emergence in 2019) as inputs (Pelech et al., 2023). Diffuse radiation was estimated using 72 direction light sources repeatedly positioned in a hemisphere in six circles each with 12 light sources. This diffuse radiation dome was randomly rotated at each time step to minimise spatial variation in light distribution. The light absorption of individual leaf organs was calculated based on the amount of light intercepted and its optical properties. Leaf reflectance was set to 0.1, and transmittance was set to 0.03 for both species (McCree, 1972). Internodes, sheaths, lateral maize ear, axillary pods, petiole and petiolules did not transmit any light, and their reflectance was set as the sum of the transmittance and reflectance of the leaves. The radiation model was invoked once per simulation step (1 day) for computation of local light absorption by all leaf organs in the scene by reverse Monte-Carlo ray-tracing algorithm in GroIMP (Hemmerling et al., 2008). Daily solar radiation between observed and simulated values during the 2019 growing season are represented in Supplementary Figure 5.

#### 3. Photosynthesis and assimilate allocation model

A model relating leaf net carbon assimilation (A_leaf_, µmol m^-2^ s^-1^) to absorbed photon flux density (PFD) was used based on light interception, according to a non-rectangular hyperbola (Thornley, 1998),

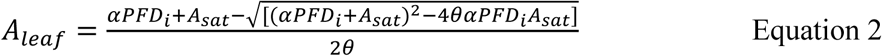

where α was the initial slope of the light response curve (apparent quantum yield), PFD_i_ (µmol m^-2^ s^-1^) was simulated light absorption by the upper leaf surface, θ was the curvature of the non-rectangular hyperbola set to 0.7, and A_sat_ (µmol m^-2^ s^-1^) was the light-saturated photosynthetic rate. Values of α and A_sat_ were derived from photosynthetic light response curves (Pelech et al., 2023) and input to the model. The simulated values of A_leaf_ were summed to calculate daily gross assimilated CO_2_ per plant per day (A_G,_ mol plant^-1^ d^-1^),

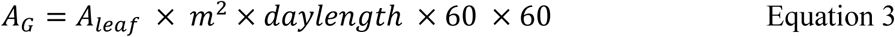

where m^2^ is the total plant leaf surface area, and daylength is the calculated hours of daylight, multiplied by the number of minutes and seconds per minute for the conversion into a day.

As previously described in Evers et al., 2010, the daily net substrates available for maize and soybean growth (B_N_) based on assimilates acquired was estimated by,

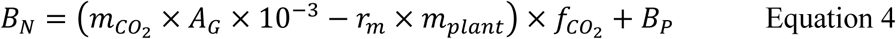

where *m*_*co*_2__ is CO_2_ molar mass (44.01, g mol^-1^ ), multiplied by 10 to convert µg to mg, and *r*_*m*_ is the fraction of total plant biomass (*m_plant_*, mg) subtracted from the number of available substrates and used by the plant for maintenance respiration per day (0.015 g CO_2 g_^-1^ plant biomass d^-1^). Variable *f*_*co*_2__ is a conversion factor (0.6) from grams of CO_2_ to grams of plant biomass, and *B_p_* is the substrate reserve pool. The available daily net substrates (B_N_) were then distributed among all growing organs: leaves (including the petiole and petiolules for soybean), internodes, sheaths, flowers, and the root (not morphologically simulated) using the relative sink strength principle (Heuvulink, 1996). The relative sink strength principle assumes that all plant substrates are equally available to all competing organs but determined by their growth rate,

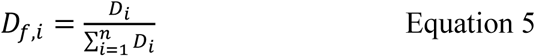

where *D_f,i_* is the relative sink strength of organ *i* (dimensionless), *D_i_* is the absolute sink strength expressed as the potential growth rate in mg of substrate organ^-1^ d^-1^, and *n* is the total number of sink organs. Values of *D_i_* were calculated according to (YIN et al., 2003/02/01), a bell-shaped curve based on the first derivative of a beta growth function.

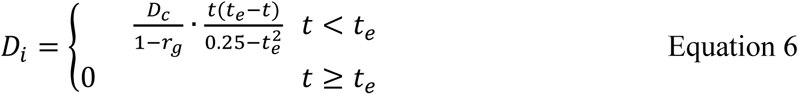

where *t_e_* is the growth duration of an organ (°C d), *t* is the amount of thermal time that passed since the start of organ growth (°C d), *r_g_* is the fraction of potential substrate lost to growth respiration (0.3), and *D_c_* is the maximum sink strength defined as,

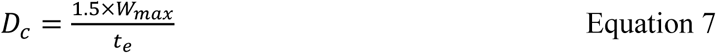

where *W_max_* is the maximum final biomass (mg organ^-1^) for leaves, internodes, flowers (ear or pods) and roots. Values of *W_max_* were input into the model derived from biomass measurements and values of *t_e_* were also input into the model derived from measurements of thermal time between organ appearance and elongation (Supplementary Table 1 and Supplementary Figure 1). The daily amount of substrate that was allocated to a sink organ (*B_i_*) was calculated as,

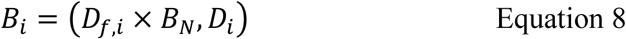

If *D_f,i_* × *B_N_* (substrates available to the organ) is higher than the sink strength (*D_i_*), the difference was added to the plant substrate reserve pool (*B_p_*, as shown in Equation 4). Values of *B_i_* were added to organ biomass daily after subtracting the potential substrate lost to growth respiration. The plant substrate reserve pool *B_p_* and biomass values of all organs were added daily to arrive at total plant mass (*m_plant_*), used in Equation 4.

### Verification simulations

Verification of model performance was compared against empirical data collected during the 2019 field experiment at the plant and field level. Verification simulations consisted of maize monoculture (M_maize_), soybean monoculture (M_soybean_), solar corridor maize monoculture (sc.M_maize_), and solar corridor intercrop (sc.I_maize+soybean_). Simulated plot dimensions are indicated in Table 1. Plasticity in this study was defined as the difference in trait values between cropping systems and trait inputs are given in Table 2. Since no significant differences in maize yield between the M_maize_ and sc.M_maize_ systems were found in Pelech et al., 2023, maize trait values remained the same for M_maize_ and sc.M_maize_ simulations. The root mean square error (RMSE) was calculated to assess the similarity between the observed and simulated values.

**Table 1.** Plot dimensions under an equal cloned area of 100 m^2^ for verification and trait analysis simulation scenarios^a^.

| Single-row system | Abbreviation | Plant density<br>(m<sup>2</sup>) | Relative<br>density<sup>b</sup> | Total plant<br>number | Row space<sup>b</sup><br>(m) |
| --- | --- | --- | --- | --- | --- |
| Maize monoculture | M<sub>maize</sub> | 6 | - | 60 | 0.76 |
| Solar corridor monoculture | sc.M<sub>maize</sub> | 6 | - | 60 | 1.52 |
| Soybean monoculture | M<sub>soybean</sub> | 24 | - | 240 | 0.76 |
| Solar corridor intercrop | sc.I<sub>maize</sub> | 6 | 1 | 60 | 1.52 |
|  | sc.I<sub>soybean</sub> | 12 | 0.5 | 120 | 1.52 |
| **Twin-row system** |  |  |  |  |  |
| Solar corridor intercrop | sc.I<sub>maize</sub> | 6 | 1 | 60 | 0.19 |
|  | sc.I<sub>soybean</sub> | 8 | 0.3 | 120 | 1.52 |

<sup>a</sup>Field dimensions for validation scenarios are indicated in Supplementary Table 2<sup>b</sup>Relative plant density with respect to the monoculture<sup>c</sup>Row spacings within species are presented, row spacings between species are indicated in Figure 2.

**Table 2.** Trait inputs for monoculture and intercrop simulations.

| Trait<sup>a</sup> | Unit | Abbreviation | Maize |  | Soybean |  |
| --- | --- | --- | --- | --- | --- | --- |
| | | | $M_{maize}$,<br>$sc.M_{maize}$<sup>b</sup> | $sc.I_{maize}$ | $M_{soybean}$ | $sc.I_{soybean}$ |
| Leaf mass per unit area | mg/cm<sup>2</sup> | LMA | 5.8 | 5.3 | 5 | 3.6 |
| Light saturated<br>photosynthetic rate | μmol m<sup>-2</sup> s<sup>-1</sup> | A<sub>sat</sub> | 56.2 | 48.8 | 37.7 | 29.9 |
| Apparent quantum yield | - | AQY | 0.07 | 0.06 | 0.06 | 0.06 |
| Specific internode length | mm/mg | SIL | 0.033 | 0.030 | 0.9 | 1 |

<sup>a</sup>Plasticity in LMA, A<sub>sat</sub>, AQY and SIL were defined as the difference in trait values between cropping systems
<sup>b</sup>Pelech et al., 2023 determined no significant yield differences between $sc.M_{maize}$ and $M_{maize}$ , therefore, all traits remained the same for simulations.

### Validation simulations

The yield output of maize and soybean monoculture simulations was validated against monoculture data on the same cultivars collected across multiple years at three locations in Illinois (data courtesy of the F.Below Laboratory, University of Illinois Urbana Champaign). Validation data was selected based on the weather conditions closest to the 2019 verification year (Pelech et al., 2023). For soybean, 2018 and 2019 data at a population density of 40 plants m^2^ with 0.76 m rows were used to compare against simulations of equal field dimensions. For maize, 2017 and 2018 data at multiple plant densities (8, 9 and 11 plants m^2^) and row spacings (0.76 m and 0.51 m) were also used to compare against simulations of equal field dimensions (Supplementary Table 2). Fitted temperature data for 2017 and 2018 were also used as inputs (Supplementary Figure 6). Each location was considered a replicate (n = 3) which had a different planting date. To represent this in the model, each replicate simulation had a different starting day of year (Supplementary Table 2), assuming 10 days until emergence from each planting date. All simulations ended after 130 days for maize and 113 days for soybean after yields had plateaued .

### Trait analysis simulations

All traits were evaluated in a complete factorial design to disentangle their effect on maize yield in M_maize_, sc.M_maize,_ and single-row sc.I_maize+soybean_ configurations, and 2) soybean yield in single-row sc.I_maize+soybean_ configuration only. For all intercrop simulations evaluating maize traits, soybean was set to the total intercrop phenotype as the first objective was to analyse the contribution of plastic maize traits to maize performance in a realistic sc.I_maize+soybean_ setting. For intercrop simulations evaluating soybean traits, maize was set to the improved phenotype determined from the first objective (Table 3).

**Table 3.** A list of the reference (*R*) and experimental (*E*) simulation scenarios for assessing the contributions of individual plastic maize traits.

| Maize trait | Simulation scenarios |  |  |  |  |  |  |  |
| --- | --- | --- | --- | --- | --- | --- | --- | --- |
|  | *R* | *E* | *E* | *E* | *R* | *E* | *E* | *E* |
| LMA | ● | ● | ○ | ○ | ○ | ○ | ● | ● |
| A<sub>sat</sub> + AQY | ● | ○ | ● | ○ | ○ | ● | ○ | ● |
| SIL | ● | ○ | ○ | ● | ○ | ● | ● | ○ |
| Soybean trait |  |  |  |  |  |  |  |  |
| LMA | ● | ● | ○ | ○ | ○ | ○ | ● | ● |
| A<sub>sat</sub> | ● | ○ | ● | ○ | ○ | ● | ○ | ● |
| SIL | ● | ○ | ○ | ● | ○ | ● | ● | ○ |

The relative contribution of the individual plastic traits to yield (g m^-2^) was calculated as the absolute difference in yield by resetting the single trait from its value in the intercrop to its value in the monoculture, and vice versa. The absolute yield difference was also normalised by expressing it as a percentage (Zhu et al., 2015),

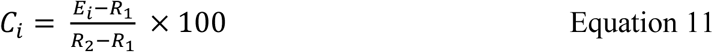

where *E_i_* is the yield of the experimental scenario with trait *i* set to either the intercrop or monoculture phenotype, *R_1_* is the yield of the reference scenario with all traits set to either the intercrop or monoculture phenotype*, R_2_* is the yield of the reference scenario with all traits set to either the intercrop or monoculture phenotype, and *C_i_* is the relative contribution of trait *i* to the difference in yield between the total intercrop and monoculture phenotype.

The yields of the single-row and twin-row arrangement of maize rows in the solar corridor intercrop (Figure 4) were evaluated as: (i) the observed total intercrop phenotype, (ii) the improved maize phenotype only, and (iii) the improved maize and soybean phenotype. Calculation of the Land Equivalent Ratio (LER) was used based on the average yield to assess the land-use efficiency across scenarios by summing the relative yields of both species. If LER > 1, the intercrop system has a yield advantage and increased land-use per unit area.

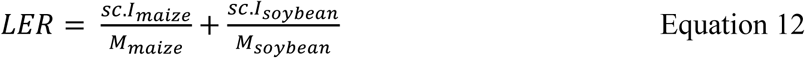

**Figure 4.**
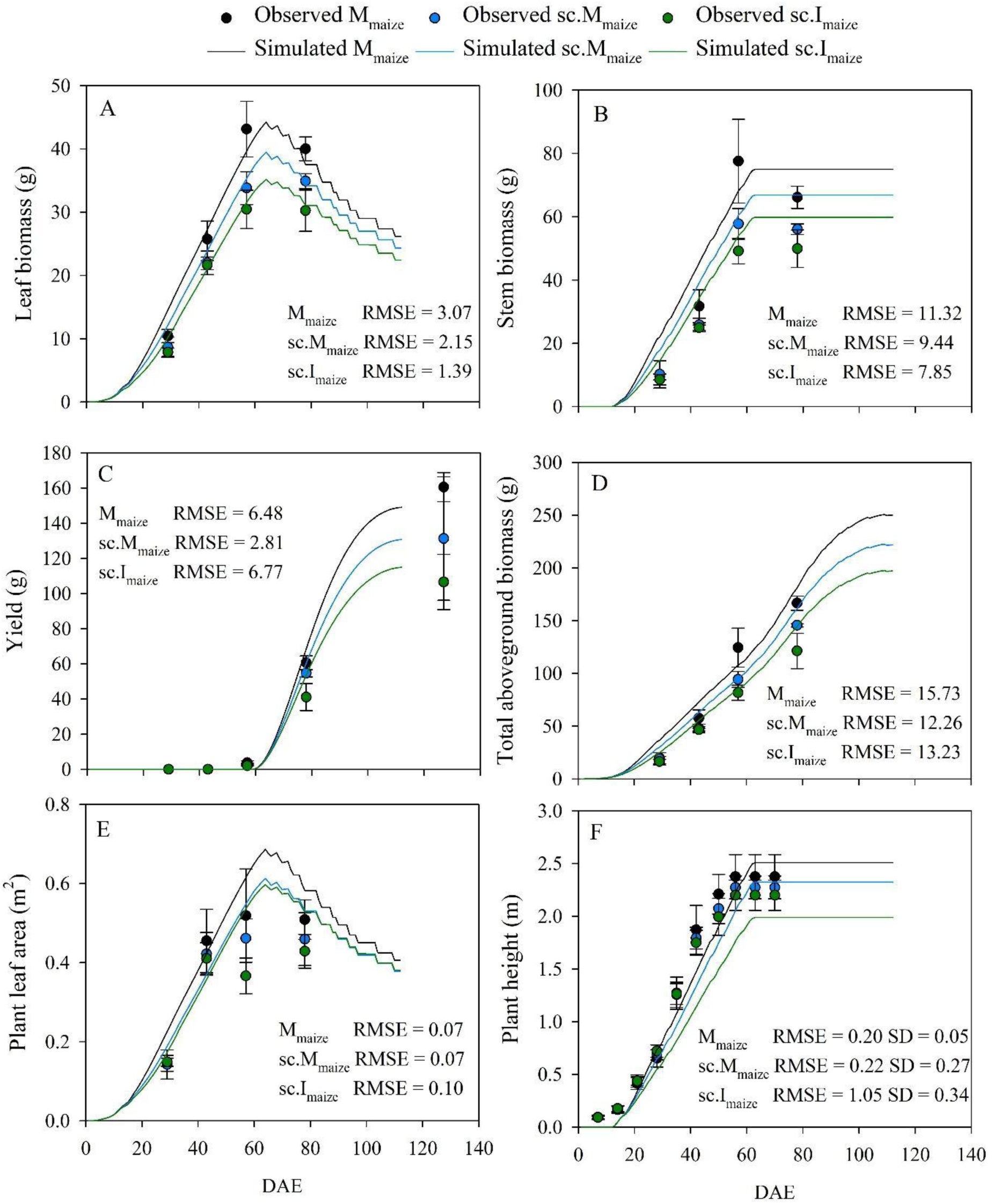
Comparisons of plant level observed (circle) and simulated (line) maize variables in monoculture (M_maize_ [black]), solar corridor monoculture (sc.M_maize_ [blue]), and solar corridor intercrop (sc.I_maize_ [green]) across days after emergence (DAE). Observed values were measured during the 2019 growing season (Pelech et al., 2023), and simulated values are the average of three maize plant simulations. Error bars represent ± SD (n = 4, 3).

### Model replication and stochastic components

All verification and trait analysis simulation scenarios were performed on an equal land area basis whereas validation scenarios were performed based on equal number of plants simulated due to the differences in plant densities and row spacings (Table 1 and Supplementary Table 2). All simulations had an internal cloned area of 10 m in both the x and y direction to minimise border effects. To ensure there were no gaps within the cloned area by differences in row lengths for the intercrop simulations, the simulated soybean plant density (24 plants m^2^) could not equal the average plant density observed in the field experiment (22 plants m^2^, Pelech *et al*., 2023) because it was not a multiple of the average maize plant density (6 plants m^2^). Therefore, verification simulations of soybean were off by two plants per m^2^ for both M_soybean_ and sc.I_maize+soybean_ configurations. Each scenario for verification, validation, and trait analysis simulations (30 scenarios in total) was replicated three times, and the average was calculated as the output variables per simulation which differed slightly due to 1) randomisation of plant orientation and 2) stochasticity inherent to the radiation model.

## Results

### Verification of model performance

The model effectively captured the architectural development of maize and soybean plants in all simulated cropping systems, including the representation of leaf ageing and subsequent senescence of individual leaves (Supplementary Figures 7-10 and Supplementary Videos 1-4).

Overall, there was similarity between the simulated and observed maize values at the plant level, excluding the simulated values of sc.I_maize_ plant height, which was slightly under-estimated relative to in-field measurements (Figure 4). Simulated light interception fractions were underestimated for all maize systems at the field level, while LAI values were overestimated (Figure 5AB). Simulated total aboveground biomass values were comparable to those measured (Figure 5C), and yield was slightly underestimated for only the M_maize_ simulations (Figure 5D).

**Figure 5.**
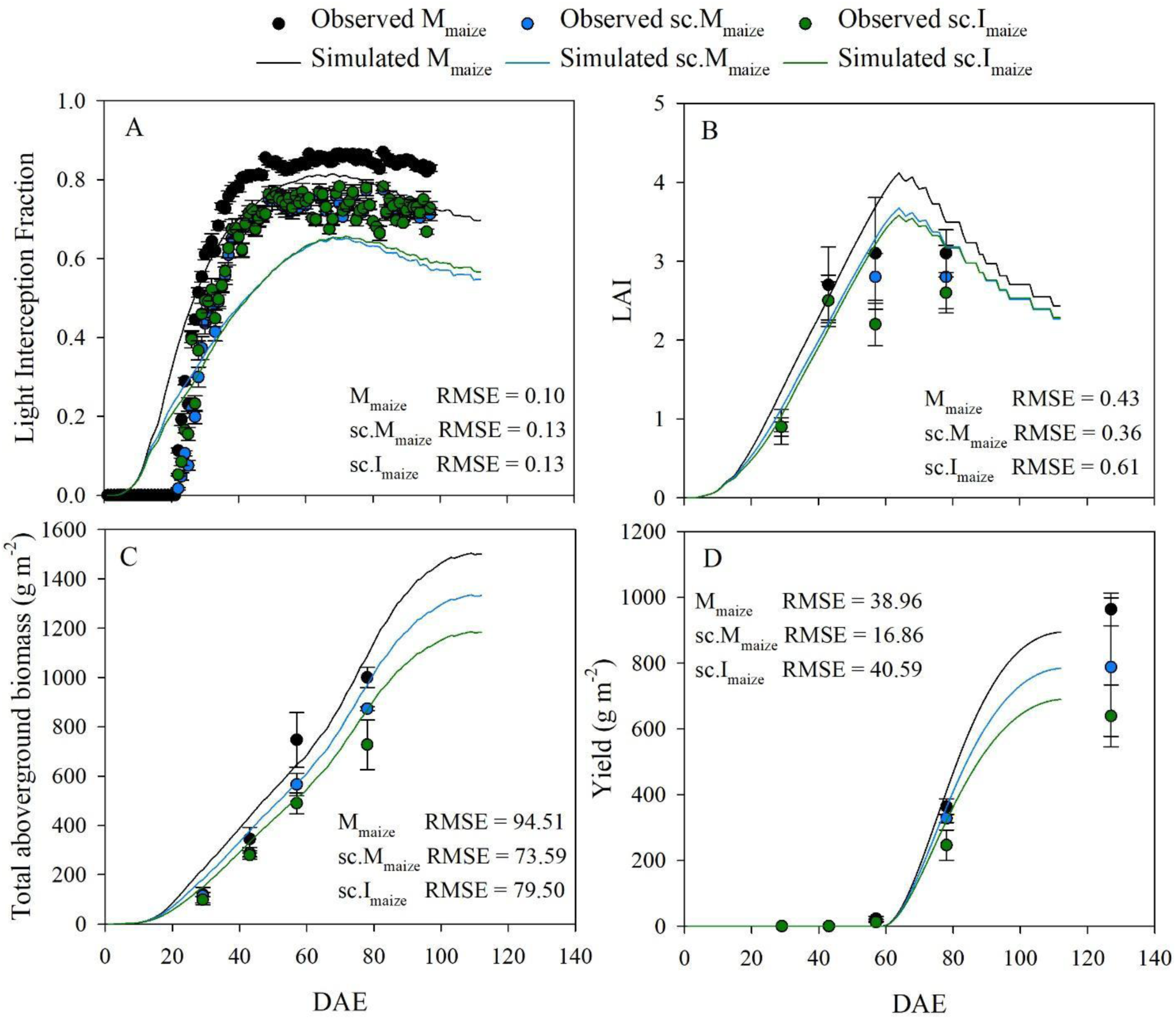
Comparisons of field level observed (circle) and simulated (line) maize variables in monoculture (M_maize_ [black]), solar corridor monoculture (sc.M_maize_ [blue]), and solar corridor intercrop (sc.I_maize_ [green]) across days after emergence (DAE). Data sources, error bars and replicates are as in Figure 3, excluding light interception fraction where error bars represent ± SE for clarity.

For individual soybeans, simulated total aboveground biomass, leaf biomass and plant height were close to measured values for sc.I_soybean_ (Figure 6ADF). Plant leaf area and yield were underestimated for simulated sc.I_soybean_ plants (Figure 6BCEF), but less so for equivalent field-level variables, LAI and yield (Figure 7). Light interception fractions were also very similar, with an RMSE of 0.04 and field-level total aboveground biomass with an RMSE of 14.03 g m^-2^ (Figure 7AC). For M_soybean_ plants, stem biomass was overestimated towards the end of the simulation, but the remaining plant level variables were within an order of magnitude with observations (Figure 6). At the field level, the RMSE for all M_soybean_ variables were higher than sc.I_soybean_, however, simulated values of LAI and yield at mature development were within the observed range (Figure 7BD).

**Figure 6.**
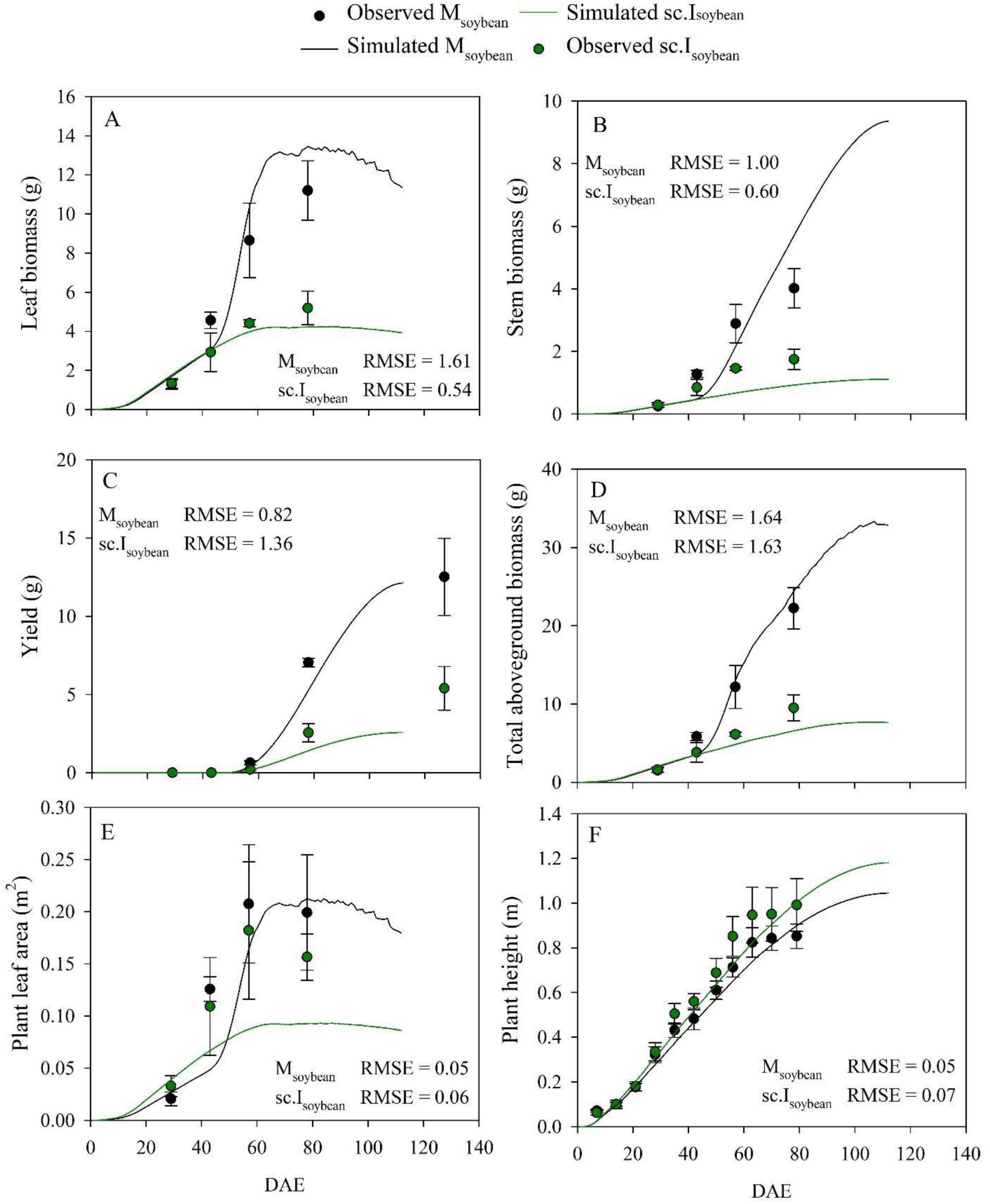
Comparisons of plant level observed (circle) and simulated (line) soybean vairables in monoculture (M_soybean_,[black]) and solar corridor intercrop (sc.I_soybean_, [green]) across days after emergence (DAE). Data sources, error bars and replicates are as in Figure 3.

**Figure 7.**
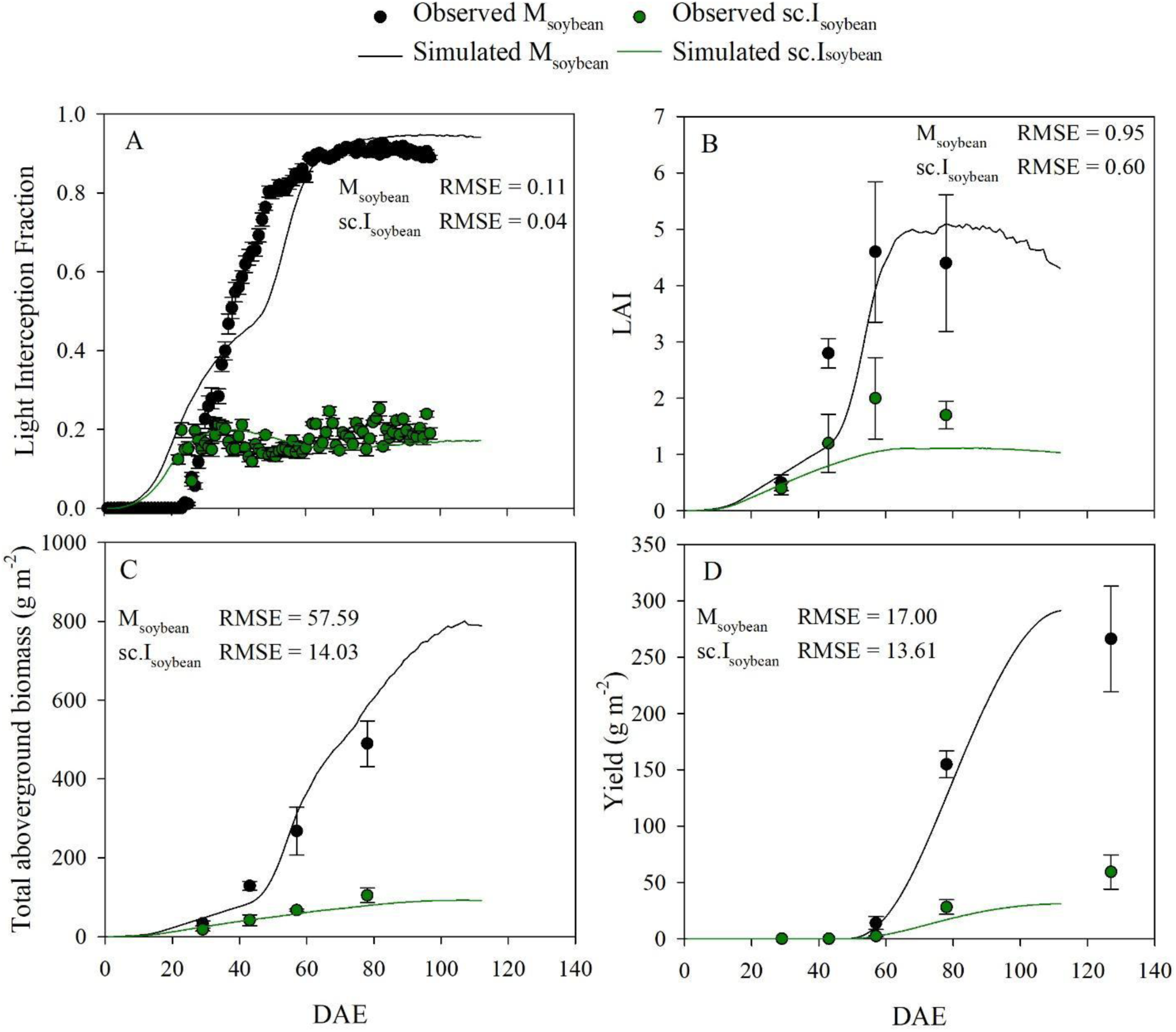
Comparisons of field level observed (circle) and simulated (line) soybean variables in monoculture (M_soybean_, [black]) and solar corridor intercrop (sc.I_soybean_, [green]) across days after emergence (DAE). Data sources, error bars and replicates are as in Figure 3.

### Validation of maize and soybean yields in monoculture configurations

Maize yield data for four monoculture configurations over two independent years was used to validate maize verification (Figure 8A). Overall, the model underestimated the observed yield between 10-27% but could mimic the directionality of the yield trends between the monoculture configurations. Soybean yield data for a single monoculture configuration over two years, including the same year as the verification experiment in this study (2019), were used to validate soybean verification (Figure 8B). Like maize, the soybean model underestimated the observed yields but was able to mimic the directionality of yield trends between the two years (Figure 8B).

**Figure 8.**
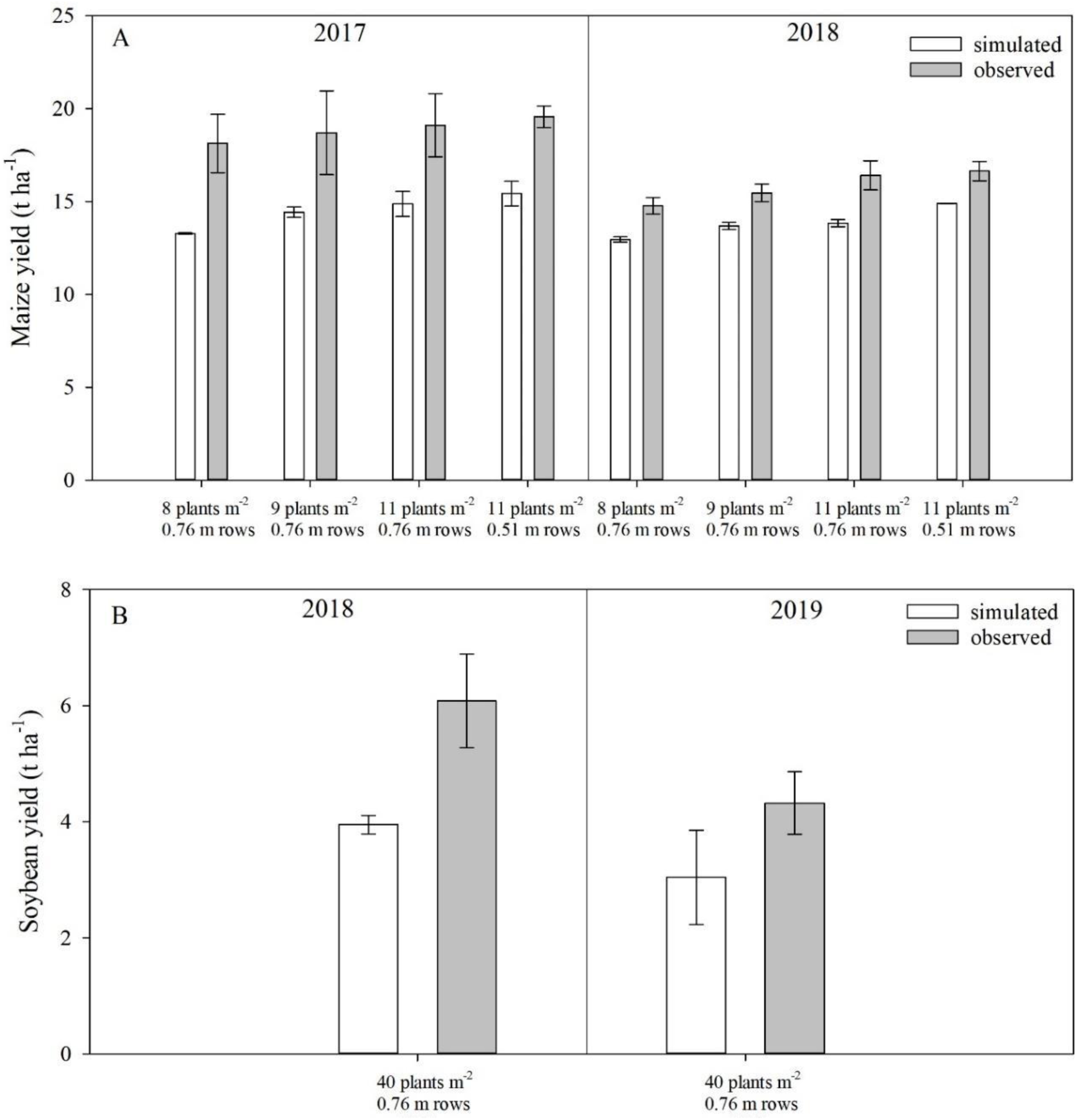
Yield validation comparisons between simulated (white bars) and observed (grey) for A maize and B soybean monocultures across different years. The corresponding plant density and row spacing are indicated on the x-axis. Observed yield values are the average from three locations in Illinois on the same cultivars used to parameterise the model. Simulated yield values are the average three independent simulations, each with 210 maize plants and 240 soybean plants. Fitted air temperature measurements were input for each year (Supplementary Figure 6), and the starting day of year was input for each replicate simulation to represent emergence at each replicate location (Supplementary Table 2). Error bars represent ± SD (n = 3). Observed data courtesy of the F.Below Laboratory.

### The relative contribution of plastic maize traits to yield

The average yield of maize with a total intercrop phenotype (all three traits with intercrop values) in the M_maize_ configuration was similar to the monoculture phenotype in the same configuration (Table 4). When the contribution of the total intercrop phenotype to maize yield in the M_maize_ configuration was partitioned into the three traits, the lower LMA value contributed +22% ± 19 SD, the lower photosynthetic rates contributed +44% ± 13 SD, and SIL contributed +22% ± 35 SD to the < 1% decrease in maize yield (Figure 9A).

**Figure 9.**
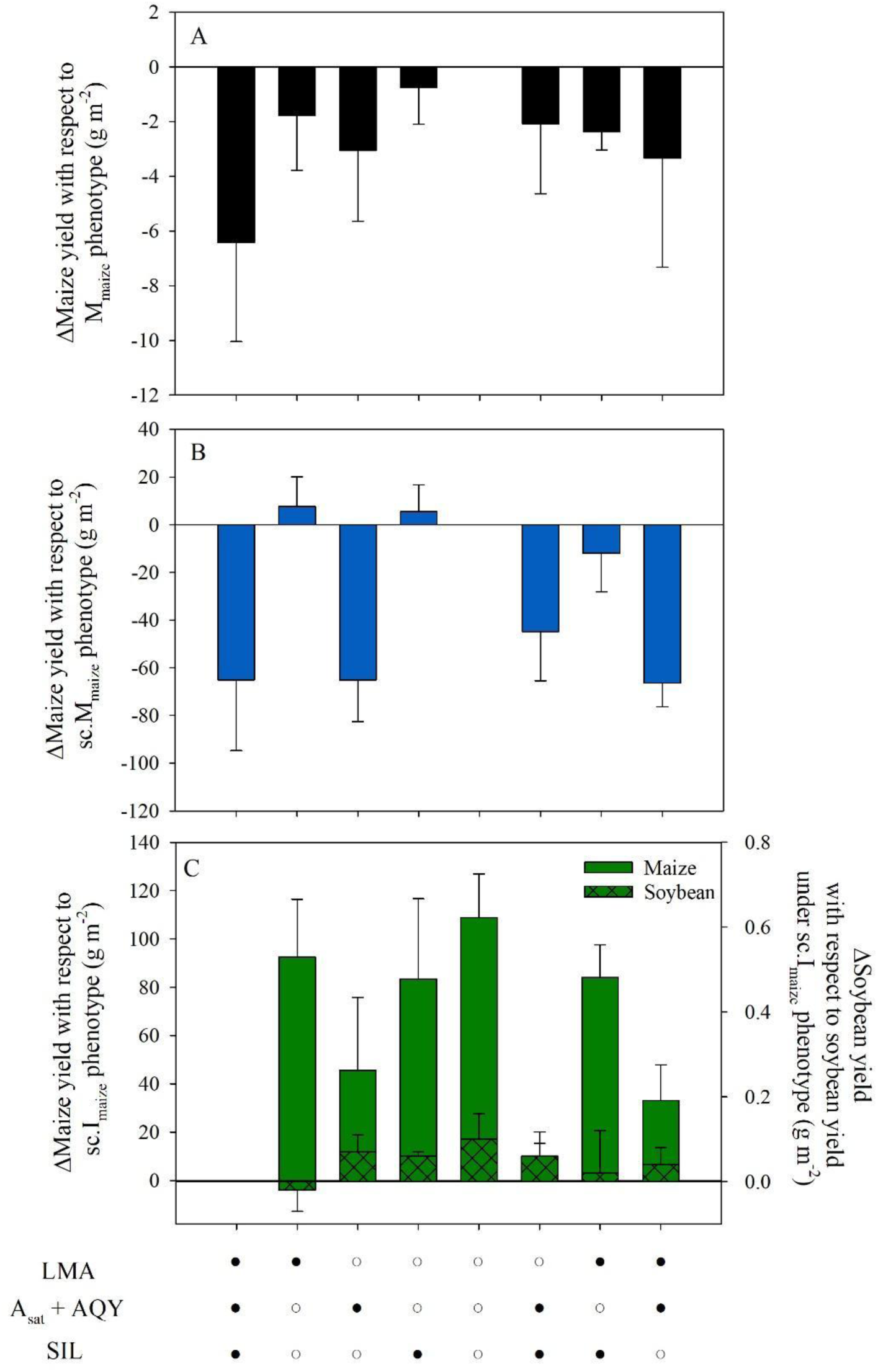
Mean maize yield differences in simulations of A maize monoculture, B maize solar corridor and C solar corridor intercrop. Black circles represent the intercrop maize trait and white circles represent the monoculture maize trait. The combination of different symbols represents the integration of maize traits. All soybean traits were set to the intercrop phenotype. Error bars represent ± SD (n=3).

**Table 4.**
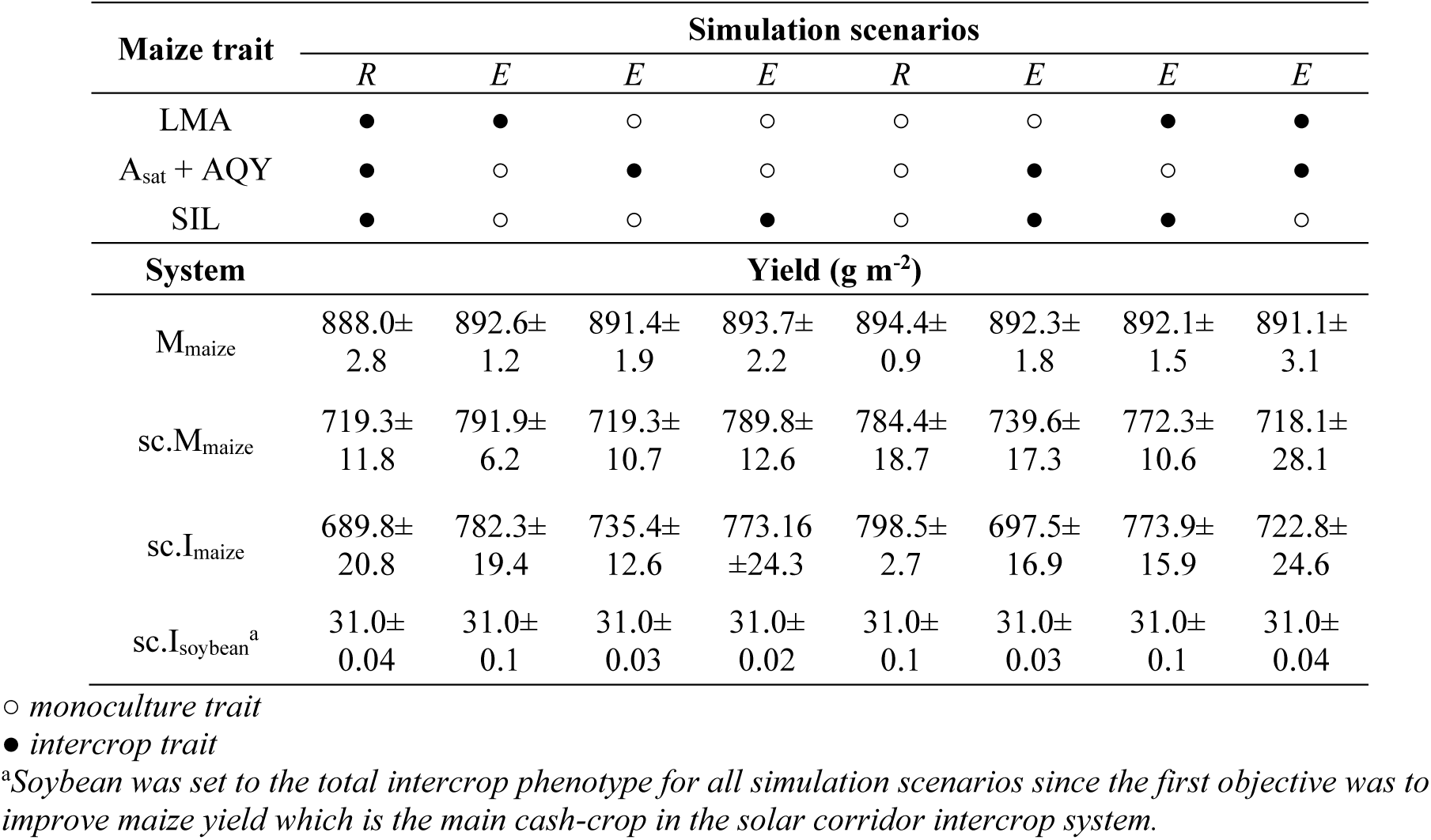
Final yield output from reference (*R*) and experimental (*E*) simulation scenarios across maize systems that disentangled the contribution of monoculture and intercrop maize traits.

| Maize trait | Simulation scenarios |  |  |  |  |  |  |  |
| --- | --- | --- | --- | --- | --- | --- | --- | --- |
|  | *R* | *E* | *E* | *E* | *R* | *E* | *E* | *E* |
| LMA | ● | ● | ○ | ○ | ○ | ○ | ● | ● |
| A<sub>sat</sub> + AQY | ● | ○ | ● | ○ | ○ | ● | ○ | ● |
| SIL | ● | ○ | ○ | ● | ○ | ● | ● | ○ |
| System | Yield (g m<sup>-2</sup>) |  |  |  |  |  |  |  |
| M<sub>maize</sub> | 888.0±<br>2.8 | 892.6±<br>1.2 | 891.4±<br>1.9 | 893.7±<br>2.2 | 894.4±<br>0.9 | 892.3±<br>1.8 | 892.1±<br>1.5 | 891.1±<br>3.1 |
| sc.M<sub>maize</sub> | 719.3±<br>11.8 | 791.9±<br>6.2 | 719.3±<br>10.7 | 789.8±<br>12.6 | 784.4±<br>18.7 | 739.6±<br>17.3 | 772.3±<br>10.6 | 718.1±<br>28.1 |
| sc.I<sub>maize</sub> | 689.8±<br>20.8 | 782.3±<br>19.4 | 735.4±<br>12.6 | 773.16<br>±24.3 | 798.5±<br>2.7 | 697.5±<br>16.9 | 773.9±<br>15.9 | 722.8±<br>24.6 |
| sc.I<sub>soybean</sub><sup>a</sup> | 31.0±<br>0.04 | 31.0±<br>0.1 | 31.0±<br>0.03 | 31.0±<br>0.02 | 31.0±<br>0.1 | 31.0±<br>0.03 | 31.0±<br>0.1 | 31.0±<br>0.04 |

○ monoculture trait
● intercrop trait
<sup>a</sup>Soybean was set to the total intercrop phenotype for all simulation scenarios since the first objective was to improve maize yield which is the main cash-crop in the solar corridor intercrop system.

Compared to the M_maize_ configuration, the contribution of individual traits to maize yield in the sc.M_maize_ configuration were of greater magnitude (Figure 9B). Maize with a total intercrop phenotype yielded 719.3 g m^-2^, equal to an 8.3% reduction compared to the maize yield associated with a total monoculture phenotype at 784.4 g m^-2^. When the contribution of the total intercrop phenotype to maize yield in the sc.M configuration was partitioned into the three traits, the lower LMA value contributed -18% ± 22 SD, the lower photosynthetic rates contributed +107% ± 35 SD, and SIL contributed -12% ± 22 SD to the 8.3% decrease in maize yield (Figure 8B). Therefore, the intercrop LMA trait value increased maize yield compared to the total monoculture phenotype, by ∼1%, and was the greatest yield achieved in the sc.M_maize_ configuration at 791.9 g m^-2^ (Table 4).

In the single-row sc.I_maize+soybean_ configuration, maize yield with a total monoculture phenotype was 798.5 g m^-2^, equal to a 14% yield gain compared to the total intercrop phenotype at 689.8 g m^-2,^ which did not cause a yield penalty to intercropped soybean (Table 4).

Ultimately, all intercrop simulation scenarios that assessed the contribution of monoculture and intercrop maize traits had no influence on soybean yield. When the contribution of the total monoculture phenotype to maize yield in the single-row sc.I_maize+soybean_ configuration was partitioned into the three traits, the higher LMA value contributed +6% ± 10 SD, the higher photosynthetic rates contributed +78% ± 11 SD, and SIL contributed +31% ± 13 SD to the 14% increase in maize yield. Furthermore, maize with a total monoculture phenotype achieved a greater yield by 2% in the single-row sc.I_maize+soybean_ configuration than in the sc.M_maize_ configuration.

### The relative contribution of soybean traits to yield in the intercrop system

The contribution of soybean traits to yield was assessed in the single-row sc.I_maize+soybean_ configuration only (Table 5 and Figure 10). When maize was set to the phenotype that gave the highest yield, the monoculture phenotype, there was no significant impact on intercropped soybean yield (Table 4 and Figure 9). Ultimately, the variations in soybean phenotype had no influence on its yield but had a small influence on maize yield. Soybean yield with a total monoculture phenotype and total intercrop phenotype were equal at 31.0 g m^-2^. However, the monoculture soybean phenotype with A_sat_ set to the intercrop trait value produced the greatest maize yield at 805.0 g m^-2^, a 2% increase in maize yield from the total monoculture phenotype.

**Figure 10.**
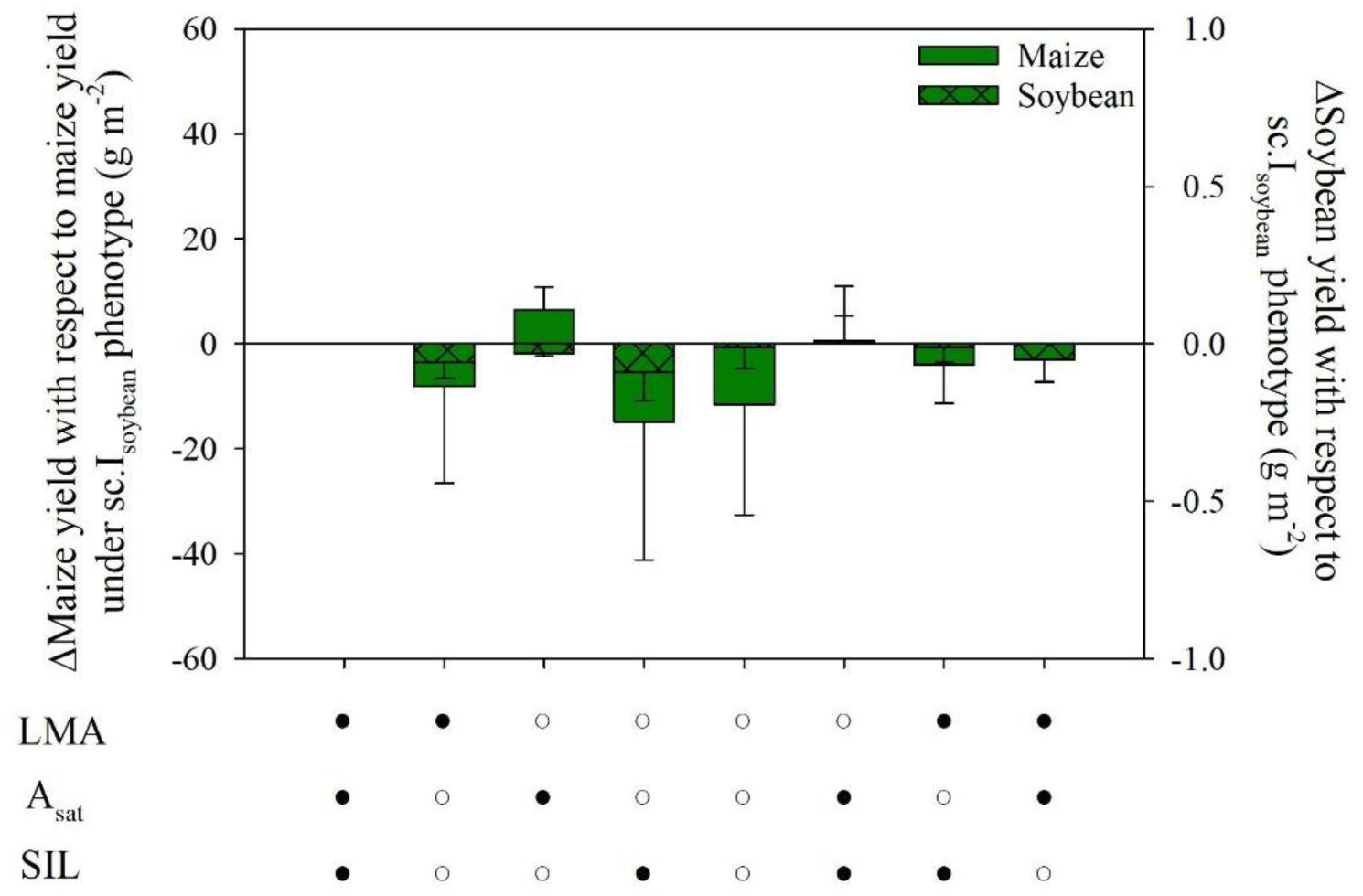
Mean soybean yield differences in solar corridor intercrop simulations. Black circles represent the intercrop soybean trait and white circles represent the monoculture soybean trait. The combination of different symbols represents the integration of soybean traits. All maize traits were set to the total monoculture phenotype. Error bars are as in Figure 8.

**Table 5.**
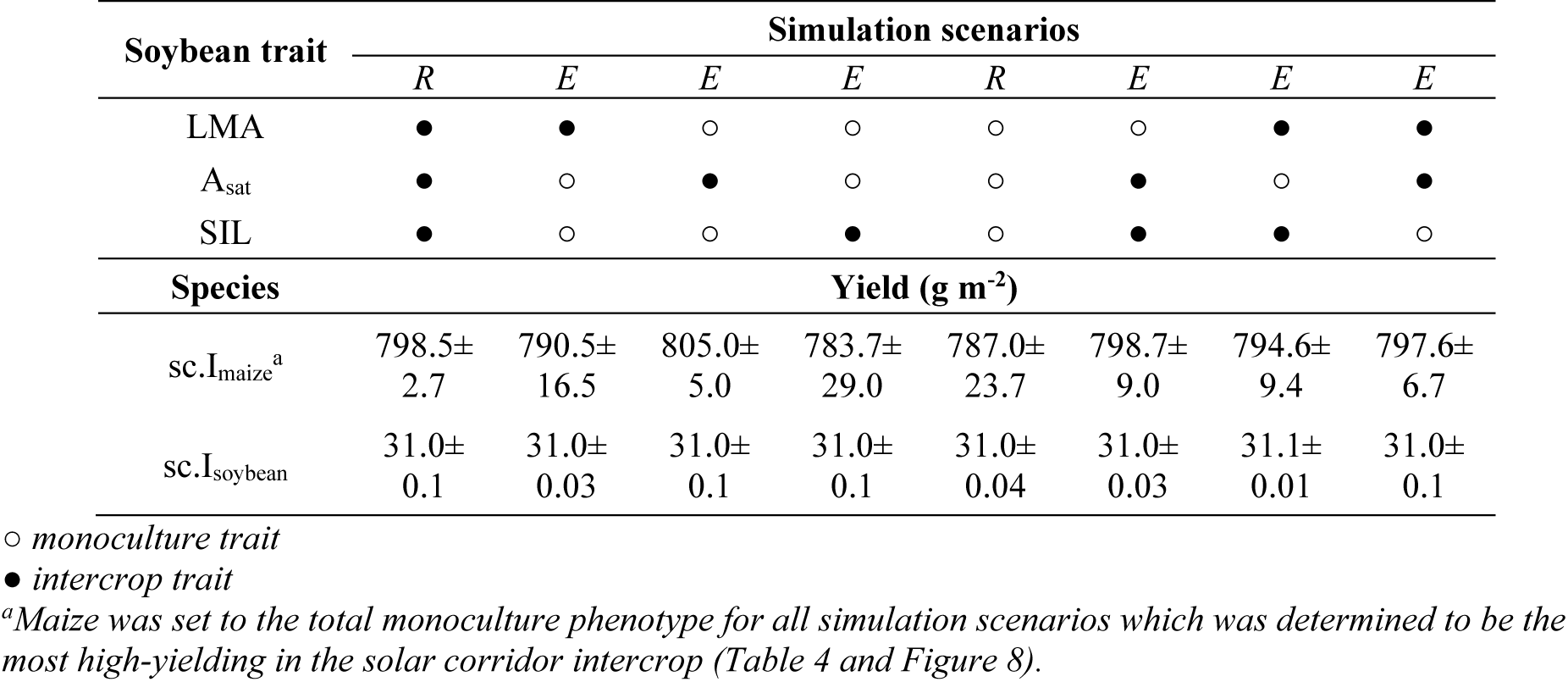
Final yield output from reference (*R*) and experimental (*E*) simulation scenarios of the solar corridor intercrop system that disentangled the contribution of monoculture and intercrop soybean traits.

### The LER and relative yield changes of the spatial arrangement of maize rows in the intercrop

The LER was calculated for simulated sc.I_maize+soybean_ systems with single-row and twin-row arrangements of maize (Table 6). The total intercrop phenotype of maize and soybean (observed) was compared against the improved maize phenotype only and the improved maize and soybean phenotype determined from the trait analysis described above. Across simulation scenarios, soybean yield in the single-row arrangement did not change but decreased by ∼33% in the twin-row arrangement. The observed scenario produced an LER of 0.88 and 0.90 for single- and twin-row sc.I_maize+soybean_ configurations, respectively. This equals a 12% and 10% decrease in land-use efficiency for the sc.I_maize+soybean_ systems compared to the standard monoculture systems of maize and soybean; however, maize yield increased by 8% in the twin-row compared to the single-row arrangement. The improved maize phenotype only produced an LER for both row arrangements equal and close to 1.00, suggesting no yield advantage compared to the monocultures of maize and soybean. Although, compared to the observed phenotype, land-use efficiency increased by 12% and 11% for single- and twin-row systems, respectively; maize yield increased by 5% in the twin-row compared to the single-row arrangement. Both improved maize and soybean phenotypes gave a slight yield advantage for both row arrangements (1-3%), but the twin-row system produced the greatest maize yield. Compared to the observed phenotype, land-use efficiency of the improved phenotypes increased by 13% for both row arrangements, and maize yield increased by 23% in the improved twin-row system compared to the observed single-row system.

**Table 6.**
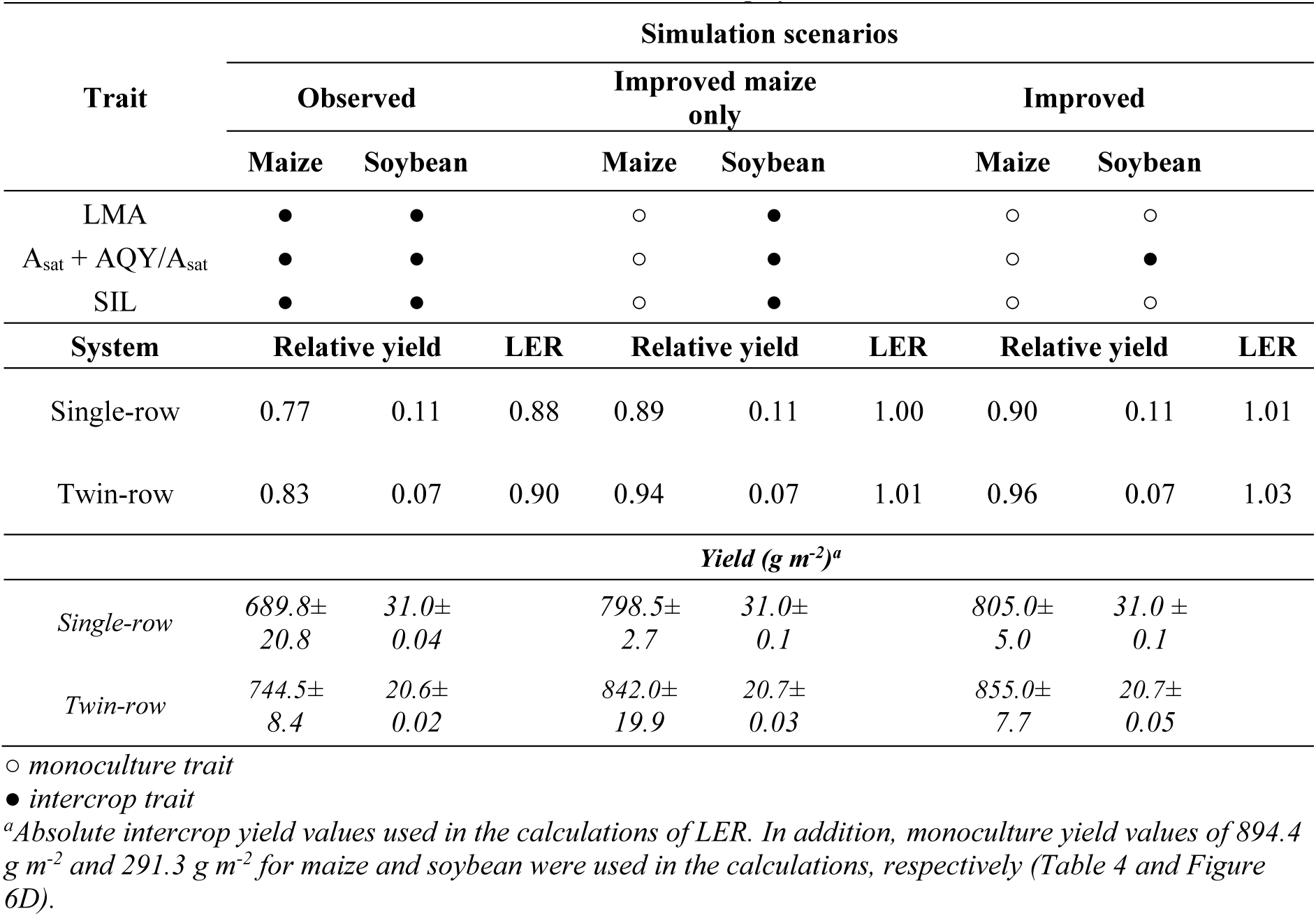
The calculation of the land equivalent ratio (LER) for observed and improved phenotypes in single- and twin-row simulations of maize in the solar corridor intercrop system.

## Discussion

This study parametrised a mechanistic maize and soybean FSP model with a representation of light interception, photosynthesis, assimilate allocation, three-dimensional plant growth and yield using data from a field experiment (Pelech et al., 2023). The model was used to identify improved maize and soybean phenotypes for the simultaneous solar corridor intercrop system by disentangling the contribution of three plastic traits to yield: photosynthesis (light-saturated rate [A_sat_] and apparent quantum yield [AQY]), leaf thickness (leaf mass per unit area [LMA]) and plant height (specific internode length [SIL]). We found that the total maize monoculture phenotype produced a 14% maize yield gain compared to the total intercrop phenotype in the single-row intercrop configuration where higher photosynthetic rates contributed the most, and greater LMA contributed the least; soybean yield was unaffected (objective 1 [Table 4 and Figure 9C]). For soybean, we discovered that the contribution of each plastic trait had no influence on soybean yield suggesting soybean plasticity is not enough to compensate for the heavy shading imposed by maize to improve soybean yields, but the monoculture soybean phenotype with a lower A_sat_ value produced a 2% maize yield gain compared to the total soybean monoculture phenotype (objective 2 [Table 5 and Figure 10]).

Finally, compared to the observed phenotypes, the improved phenotypes discovered here increased land-use efficiency by 12% and 11% for single- and twin-row systems, respectively (objective 3 [Table 6]). Ultimately, the simultaneous solar corridor intercrop system within the physiological capacity of the particular genotypes explored here can potentially increase land-use efficiency between 1-3% compared to the standard monoculture systems of the Midwest USA.

### The spatial configuration and phenotypic plasticity of maize in the solar corridor

This is the first comprehensive study to explore the role of maize traits in driving yields in a simultaneous and additive maize and soybean intercrop. The FSP modelling studies on intercrops involving maize have focused on the plasticity of the subordinate species and its role in establishing complementary light capture (Zhu et al., 2015; Li et al., 2021). The solar corridor system, however, involves the rearrangement of maize plants into wider but denser rows to maintain equal plant density which has been shown to be limiting to maize performance compared to the standard M_maize_ system (Nelson, 2014; Pelech et al., 2023). Therefore, it is critical to understand whether the maize traits within the physiological capacity of a particular genotype can overcome the limitations of 1) intraspecific light competition by doubling the maize plant density within the maize rows (sc.M_maize_ configuration) and 2) intra- and interspecific light competition by addition of a single soybean row between the wide and dense maize rows (single-row sc.I_maize+soybean_ configuration).

The plastic traits that were the most influential to maize productivity in all the configurations were the light-saturated photosynthetic rate (A_sat_) and the apparent quantum yield (AQY). Their contribution to maize productivity also strengthened from 44% in M_maize_ to 78-107% in the sc.M_maize_ and single-row sc.I_maize+soybean_ configurations (Table 4 & Figure 9). The strengthening of the role A_sat_ and AQY contributed to maize yield between the configurations suggests maintaining higher trait values is crucial to tolerating the intraspecific light competition between maize plants within the dense solar corridor rows as well as the addition of soybean in the intercrop. Typically, shade tolerant plants have a lower A_sat_ but maintain quantum efficiency for photosynthetic CO_2_ reduction (AQY), a competitive strategy to minimise carbon assimilation losses (Givnish, 1988; Valladares et al., 2008). This strategy was not realised in the intercrop phenotype, given by a lower AQY value (Table 2), and previous research suggests that the potential carbon gain of dense stands of C_4_ crops is limited by a decline in quantum efficiency with canopy depth (Pignon et al., 2017). Further research suggests that this maladaptive response to self-shading is due to light, not leaf age (Collison et al., 2020) and is less severe for erectophile than planophile leaves (Jaikumar et al., 2021). Therefore, the contribution of A_sat_ and AQY plasticity may be much greater than reported here, and future research should consider inputting trait values across phytomers. We also acknowledge that in Pelech et al., 2023, the reduction in A_sat_ was only statistically significant for soybean but given the significant reduction in light interception and energy conversion efficiency, investigating the plasticity in A_sat_ was an equal measure for the model.

### The role of phenotypic plasticity for soybean in the solar corridor intercrop system

The analysis of plastic soybean traits revealed no significant influence on soybean yield but the monoculture soybean phenotype with A_sat_ set to the intercrop trait value produced a 2% increase in maize yield (Table 5 & Figure 10). Yield improvements in the single and twin-row sc.I_maize+soybean_ configurations still produced very low soybean yields compared to its monoculture system at half the plant density (Table 6). Previous studies on other simultaneous maize and soybean intercrop configurations have also reported that although plastic soybean responses increased light capture, they are not enough to compensate for the heavy shading by maize (Li et al., 2020; Li et al., 2021; Liu et al., 2017).

In intercropping systems where both species are cultivated for cash-crop production, it is undesirable for the dominant species to improve its performance at the expense of the subordinate species, since both crops are intended to contribute to overall yield and profitability The solar corridor intercrop system, however, is considered a maize system while the intercropped legume can provide soil fertility or forage for grazing livestock after mechanical maize harvest (Deichman and Kremer, 2019). Therefore, the twin-row sc.I_maize+soybean_ configuration may be better suited for farmers based on the smaller yield penalty in maize with improved phenotypes compared to the single-row sc.I_maize+soybean_ configuration (Table 6).

### Model assumptions and limitations

For simplicity and conservation of computational power, the model assumes that the source/sink ratio of plant organs, photosynthesis, leaf optical properties and the maximum organ size and mass are constant along the plant structure (i.e. phytomers) and do not change during development. Leaf optical properties were also constant between maize and soybean, however, they differ slightly in real plants and vary based on leaf N content (McCree, 1972). Other factors that impact plant performance, such as plant N economy, water interactions and vulnerability to mechanical damage (lodging), were also not considered as they were beyond the scope of this study. Thus yield differences are driven by the impact of the intercrop configuration on the amount of light intercepted, and CO_2_ assimilated.

The amount of assimilate available for growth at each time point (source capacity, W_max_), and the potential growth rate under non-limiting assimilate supply (sink strength, t_e_) were described for each type of plant organ, including the root organ, even though roots were not simulated morphologically. This carbon allocation model also assumes that assimilates for growth are equally available to all competing sinks. While the availability of assimilate is likely involved in the magnitude of the responses, it is not the primary stimulus. Additional light cues can trigger and strengthen these shade avoidance strategies, such as a decrease in red:far-red ratio, blue light, and light intensity (Franklin, 2008; Gommers et al., 2013; Pierik & de Wit, 2014). In contrast, the plasticity in this study was defined as the difference in trait values between monoculture and intercropping systems which were constant across phytomers and developmental stages. It was also acknowledged that the experimental field study only considered light competition, which may not have been the original driver of competition to cause the trait differences between cropping systems (Pelech et al., 2023). Therefore, this study also assumes that trait differences were caused by light competition only.

Overall, the model predicted the observed differences in growth and yield between the cropping systems with different spatial configurations well at the plant and field level (Figures 4 – 7). Compared to maize, soybean inherently has more individual organs increasing the sources of error and computational power requirements; the 240 individual soybean plant simulations in the M_soybean_ configuration had over 100,000 plant organs in the scene. Therefore, the model matched direct observations better for maize than soybean. The model was also validated against yield data for the same maize and soybean cultivars in different monoculture configurations and across multiple years. All validation scenarios underestimated the average yield observed, but the model could mimic the directionality of the increasing yield trends (Figure 7). Differences in N fertiliser management between the verification experiment (202 kg N ha^-1^) and the different locations that produced the validation dataset (∼270 kg N ha^-1^) may explain why the model underestimated the observed yields.

## Conclusions

A mechanistic FSP model explored the potential to improve yields in a simultaneous solar corridor intercrop by disentangling the contribution of three plastic traits related to photosynthesis, leaf thickness and plant heigh, and tested the improved phenotypes in two intercrop configurations: single- and twin-rows of maize. The study revealed that for maize, photosynthesis had the greatest contribution (+78%), followed by specific internode length (+31%) and leaf mass per unit area (+6%) where the total maize monoculture phenotype produced the greatest maize yield without affecting the yield of intercropped soybean. Soybean trait plasticity had a negligible effect on soybean yield, but the soybean monoculture phenotype with a low photosynthetic rate resulted in the greatest intercropped maize yield. This suggests soybean plasticity is not enough to compensate for the heavy shading imposed by maize to improve soybean yields. And finally, with the improved phenotypes identified in this study, land-use efficiency can potentially increase by 1-3% compared to the standard monoculture systems of the Midwest, USA. These results can aid the selection of maize and soybean germplasm that could improve the productivity of a simultaneous solar corridor intercrop.

## Acknowledgments

We thank Stephen Long for his recommendations with drafting the manuscript. This work was funded by the Global Change and Photosynthesis Research Unit of the U.S. Department of Agriculture (USDA) Agricultural Research Service. Mention of trade names or commercial products in this publication is solely to provide specific information and does not imply recommendation or endorsement by the USDA. USDA is an equal opportunity provider and employer. Any opinions, findings, and conclusions or recommendations expressed in this publication are those of the author(s) and do not necessarily reflect the views of the USDA.

## References

Barillot, R., Escobar-Gutiérrez, J. A., Fournier, C., Huynh, P., & Combes, D. (2014). Assessing the effects of architectural variations on light partitioning within virtual wheat–pea mixtures. Annals of Botany, 114(4), 725–737. 10.1093/aob/mcu099

Bongers, J. F., Evers, B. J., & Anten, R. P. N. (2025). Plastic responses for intercrop functioning. npj Sustainable Agriculture, 3(1). 10.1038/s44264-025-00048-2

Collison, F. R., Raven, C. E., Pignon, P. C., & Long, P. S. (2020). Light, Not Age, Underlies the Maladaptation of Maize and Miscanthus Photosynthesis to Self-Shading. Frontiers in Plant Science, 11. 10.3389/fpls.2020.00783

Deichman, C. L. (2000). U.S. Patent No. 6,052,941. Washington, DC: U.S. Patent and Trademark Office.

Deichman, C. L., & Kremer, R. J. (Eds.). (2019). The Solar Corridor Crop System: Implementation and Impacts. Academic Press.

Elhakeem, A., Werf, D. V. W., Ajal, J., Lucà, D., Claus, S., Vico, A. R., & Bastiaans, L. (2019).Cover crop mixtures result in a positive net biodiversity effect irrespective of seeding configuration. Agriculture, Ecosystems & Environment, 285, 106627. 10.1016/j.agee.2019.106627

Evers, B. J., & Bastiaans, L. (2016). Quantifying the effect of crop spatial arrangement on weed suppression using functional-structural plant modelling. Journal of Plant Research, 129(3), 339–351. 10.1007/s10265-016-0807-2

Evers, B. J., Letort, V., Renton, M., & Kang, M. (2018). Computational botany: advancing plant science through functional–structural plant modelling. Annals of Botany, 121(5), 767–772. 10.1093/aob/mcy050

Evers, B. J., Werf, D. V. W., Stomph, J. T., Bastiaans, L., & Anten, R. P. N. (2019). Understanding and optimizing species mixtures using functional–structural plant modelling. Journal of Experimental Botany, 70(9), 2381–2388. 10.1093/jxb/ery288

Evers, J. B., Vos, J., Yin, X., Romero, P., van der Putten, P. E. L., & Struik, P. C. (2010). Simulation of wheat growth and development based on organ-level photosynthesis and assimilate allocation. Journal of Experimental Botany, 61(8), 2203–2216. 10.1093/jxb/erq025

Franklin, K. A. (2008/09/01). Shade avoidance. New Phytologist, 179(4). 10.1111/j.1469-8137.2008.02507.x

Gaudio, N., Escobar-Gutiérrez, J. A., Casadebaig, P., Evers, B. J., Gérard, F., Louarn, G., Colbach, N., Munz, S., Launay, M., Marrou, H., Barillot, R., Hinsinger, P., Bergez, J.-E., Combes, D., Durand, J.-L., Frak, E., Pagès, L., Pradal, C., Saint-Jean, S.,…Justes, E. (2019). Current knowledge and future research opportunities for modeling annual crop mixtures. A review. Agronomy for Sustainable Development, 39(2). 10.1007/s13593-019-0562-6

Givnish, T. (1988). Adaptation to Sun and Shade: a Whole-Plant Perspective. Functional Plant Biology, 15(2). 10.1071/pp9880063

Gommers, C. M. M., Visser, E. J. W., Onge, K. R. S., Voesenek, L. A. C. J., & Pierik, R. (2013). Shade tolerance: when growing tall is not an option. Trends in Plant Science, 18(2). 10.1016/j.tplants.2012.09.008

Hemmerling, R., Kniemeyer, O., Lanwert, D., Kurth, W., & Buck-Sorlin, G. (2008). The rule-based language XL and the modelling environment GroIMP illustrated with simulated tree competition. Functional Plant Biology, 35(10), 739. 10.1071/FP08052

Heuvelink, E. (1996). Dry Matter Partitioning in Tomato: Validation of a Dynamic Simulation Model. Annals of Botany, 77(1). 10.1006/anbo.1996.0009

Jaikumar, S. N., Stutz, S. S., Fernandes, B. S., Leakey, B. D. A., Bernacchi, J. C., Brown, J. P., & Long, P. S. (2021). Can improved canopy light transmission ameliorate loss of photosynthetic efficiency in the shade? An investigation of natural variation in *Sorghum bicolor*. Journal of Experimental Botany, 72(13), 4965–4980. 10.1093/jxb/erab176

Kremer, R. J., & Deichman, L. R. C. (2016). The solar corridor: a new paradigm for sustainable crop production. Advances in Plants & Agriculture Research, 4, 273–274.

Li, S., Evers, B. J., Werf, D. V. W., Wang, R., Xu, Z., Guo, Y., Li, B., & Ma, Y. (2020). Plant architectural responses in simultaneous maize/soybean strip intercropping do not lead to a yield advantage. Annals of Applied Biology, 177(2), 195–210. 10.1111/aab.12610

Li, S., Werf, D. V. W., Zhu, J., Guo, Y., Li, B., Ma, Y., & Evers, B. J. (2021). Estimating the contribution of plant traits to light partitioning in simultaneous maize/soybean intercropping. Journal of Experimental Botany, 72(10), 3630–3646. 10.1093/jxb/erab077

Liu, X., Rahman, T., Song, C., Su, B., Yang, F., Yong, T., Wu, Y., Zhang, C., & Yang, W. (2017/01/01). Changes in light environment, morphology, growth and yield of soybean in maize-soybean intercropping systems. Field Crops Research, 200. 10.1016/j.fcr.2016.10.003

Louarn, G., & Song, Y. (2020). Two decades of functional–structural plant modelling: now addressing fundamental questions in systems biology and predictive ecology. Annals of Botany, 126(4), 501–509. 10.1093/aob/mcaa143

McCree, K. J. (1972). The action spectrum, absorptance and quantum yield of photosynthesis in crop plants. Agricultural Meteorology, 9. 10.1016/0002-1571(71)90022-7

Nelson, K. A. (2014). Corn Yield Response to the Solar Corridor in Upstate Missouri. Agronomy Journal, 106(5). 10.2134/agronj2012.0326

Pelech, A. E., Evers, B. J., Pederson, L. T., Drag, W. D., Fu, P., & Bernacchi, J. C. (2023). Leaf, plant, to canopy: A mechanistic study on aboveground plasticity and plant density within a maize–soybean intercrop system for the Midwest, USA. Plant, Cell & Environment, 46(2), 405–421. 10.1111/pce.14487

Pierik, R., & de Wit, M. (2014). Shade avoidance: phytochrome signalling and other aboveground neighbour detection cues. Journal of Experimental Botany, 65(11). 10.1093/jxb/ert389

Pignon, P. C., Jaiswal, D., Mcgrath, M. J., & Long, P. S. (2017). Loss of photosynthetic efficiency in the shade. An Achilles heel for the dense modern stands of our most productive C 4crops? Journal of Experimental Botany, 68(2), 335–345. 10.1093/jxb/erw456

Postma, A. J., & Lynch, P. J. (2012). Complementarity in root architecture for nutrient uptake in ancient maize/bean and maize/bean/squash polycultures. Annals of Botany, 110(2), 521–534. 10.1093/aob/mcs082

Schneider, M. H. (2022). Characterization, costs, cues and future perspectives of phenotypic plasticity. Annals of Botany, 130(2), 131–148. 10.1093/aob/mcac087

Sultan, S. E. (2000). Phenotypic plasticity for plant development, function and life history. Trends in Plant Science, 5(12), 537–542. 10.1016/s1360-1385(00)01797-0

Thornley, J. H. M. (1998). Dynamic Model of Leaf Photosynthesis with Acclimation to Light and Nitrogen. Annals of Botany, 81(3), 421–430. 10.1006/anbo.1997.0575

Tilman, D., & Snell-Rood, E. C. (2014). Diversity breeds complementarity. Nature, 515(7525), 44–45. 10.1038/nature13929

Valladares, F., Gianoli, E., & Gómez, M. J. (2007). Ecological limits to plant phenotypic plasticity. New Phytologist, 176(4), 749–763. 10.1111/j.1469-8137.2007.02275.x

Valladares, F., Niinemets, Ü., Valladares, F., & Niinemets, Ü. (2008). Shade Tolerance, a Key Plant Feature of Complex Nature and Consequences. Annual Review of Ecology, Evolution, and Systematics, 39(Volume 39, 2008). 10.1146/annurev.ecolsys.39.110707.173506

Vos, J., Evers, B. J., Buck-Sorlin, H. G., Andrieu, B., Chelle, M., & Visser, D. B. H. P. (2010). Functional–structural plant modelling: a new versatile tool in crop science. Journal of Experimental Botany, 61(8), 2101–2115. 10.1093/jxb/erp345

Yin, X., Goudriaan, J., Lantinga, E. A., Vos, J., & Spiertz, H. J. (2003/02/01). A Flexible Sigmoid Function of Determinate Growth. Annals of Botany, 91(3). 10.1093/aob/mcg029

Zhu, J., Werf, D. V. W., Anten, R. P. N., Vos, J., & Evers, B. J. (2015). The contribution of phenotypic plasticity to complementary light capture in plant mixtures. New Phytologist, 207(4), 1213–1222. 10.1111/nph.13416

