## Supplementary Figure for "Disentangling the contribution of trait plasticity to improve the productivity of a maize-soybean intercrop system for the Midwest, USA"

**Supplementary Table 1 | Plant model input parameters for maize and soybean.**

| Parameter | Unit | Maize | Soybean | Source |
|---|---|---|---|---|
| Harvest time after emergence | days | 109 | 109 | Pelech *et al.*, 2023 |
| Plastochron | °C d | 47 | 57.9 | Supplementary Figure 1 |
| Final phytomer number | - | 17 | na | Pelech *et al.*, 2023 |
| Max root biomass ($W_{max}$) | mg | 10000 | 90 | Manual optimization |
| Max flower biomass ($W_{max}$) | mg | 166000 | 200 | Pelech *et al.*, 2023 |
| Max internode biomass ($W_{max}$) | mg | 5500 | 60 | Pelech *et al.*, 2023 |
| Max leaf biomass ($W_{max}$) | mg | 2900 | 400 | Pelech *et al.*, 2023 |
| Root growth duration ($t_e$) | °C d | 1800 | 180 | Manual optimization |
| Flower growth duration ($t_e$) | °C d | 500 | 690 | Manual optimization |
| Internode growth duration ($t_e$) | °C d | 60 | 90 | Manual optimization |
| Leaf growth duration ($t_e$) | °C d | 250 | 133 | Supplementary Figure 2 |
| Max internode width | m | 0.06 | 0.02 | Manual optimization |
| Specific internode length (SIL) | mm/mg | 0.033 | 0.9 | Pelech *et al.*, 2023 |
| Light saturated photosynthetic rate ($A_{sat}$)[a] | μmol $m^{-2}$ $s^{-1}$ | 56.2 | 37.7 | Pelech *et al.*, 2023 |
| Apparent quantum yield (AQY)[a] | - | 0.07 | 0.06 | Pelech *et al.*, 2023 |
| Leaf mass per unit area (LMA)[a] | mg/$cm^2$ | 5.8 | 5 | Pelech *et al.*, 2023 |
| Leaf life | - | 3.5 | 7 | Supplementary Figure 2 |
| Leaf length to width ratio | - | 7 | 2 | Supplementary Figure 3 |
| Max leaf width location | - | 0.6235 | 0.5 | Manual optimization |
| Leaf insertion angle | ° | 30 | 45 | Supplementary Figure 4 |
| Leaf curvature | ° | 120 | 30 | Manual optimization |
| Biomass fraction to sheath | - | 0.1 | na | Manual optimization |
| Specific sheath length | mm/mg | 0.5 | na | Manual optimization |
| Biomass fraction to petiole | - | na | 0.2 | Manual optimization |
| Biomass fraction to petiolule | - | na | 0.01 | Manual optimization |
| Specific petiole length | mm/mg | na | 2.2 | Pelech *et al.*, 2023 |
| Specific petiolule length | mm/mg | na | 2 | Pelech *et al.*, 2023 |
| Phyllotaxis of lower phytomers | ° | 137 | 90 | Manual optimization |
| Phyllotaxis of upper phytomers | ° | 180 | 150 | Manual optimization |
| Seed mass | mg | 360 | 187 | Pelech *et al.*, 2023 |
| Base temperature | °C | 10 | 10 | MRCC[b] |

[a]*Values represent the monoculture phenotype*

[b]*Midwestern Regional Climate Center (MRCC), cli-MATE (https://mrcc.illinois.edu/CLIMATE/)*

**Supplementary Table 2 | Field dimensions for validation simulation scenarios.**

| System | Year | Location (IL) | Starting DOY[a] | Row space (m) | Plant density ($m^2$) | Cloned area ($m^2$) | Plants simulated | Days simulated |
|---|---|---|---|---|---|---|---|---|
| Maize | 2017 | Yorkville<br>Champaign<br>Harrisburg | 146<br>119<br>139 | 0.76 | 8<br>9<br>11 | 2625<br>2333<br>1909 | 210 | 130 |
| | | | | 0.51 | 11 | 1909 | | |
| | 2018 | Yorkville<br>Champaign<br>Harrisburg | 137<br>117<br>121 | 0.76 | 8<br>9<br>11 | 2625<br>2333<br>1909 | 210 | 130 |
| | | | | 0.51 | 11 | 1909 | | |
| Soybean | 2018 | Yorkville<br>Champaign<br>Harrisburg | 146<br>146<br>132 | 0.76 | 40 | 600 | 240 | 113 |
| | 2019 | Yorkville<br>Champaign<br>Ewing | 172<br>146<br>171 | 0.76 | 40 | 600 | 240 | 113 |

*[a]Day of year, assuming 10 days between planting and emergence*

*Data courtesy of the F.Below Laboratory, University of Illinois Urbana Champaign*

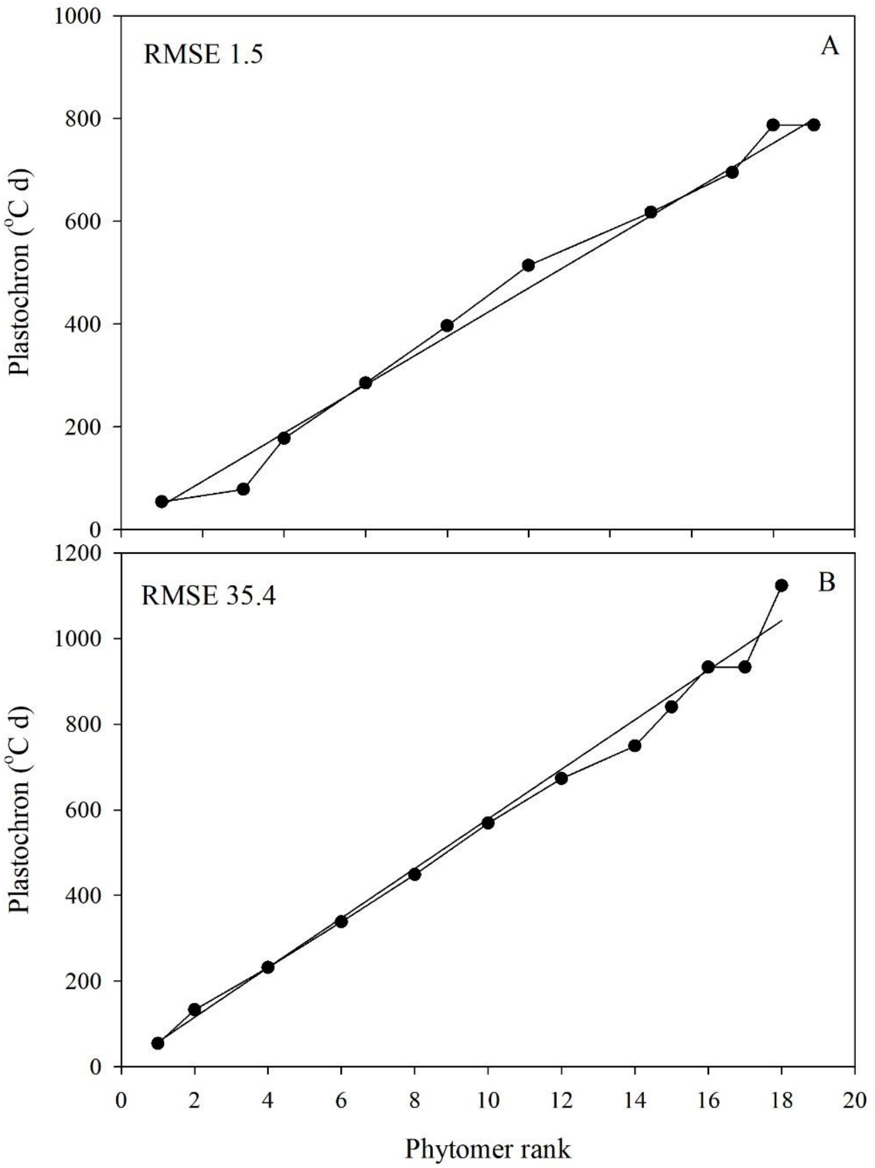


**Supplementary Figure 1.** Cumulative thermal time of fully developed phytomers across ranks in monoculture maize (A) and monoculture soybean (B). The slope of the fitted regression line is equal to the average thermal time between the creation of two phytomers (plastochron) assuming a base temperature of 10°C for both species. The plastochron was an input parameter.

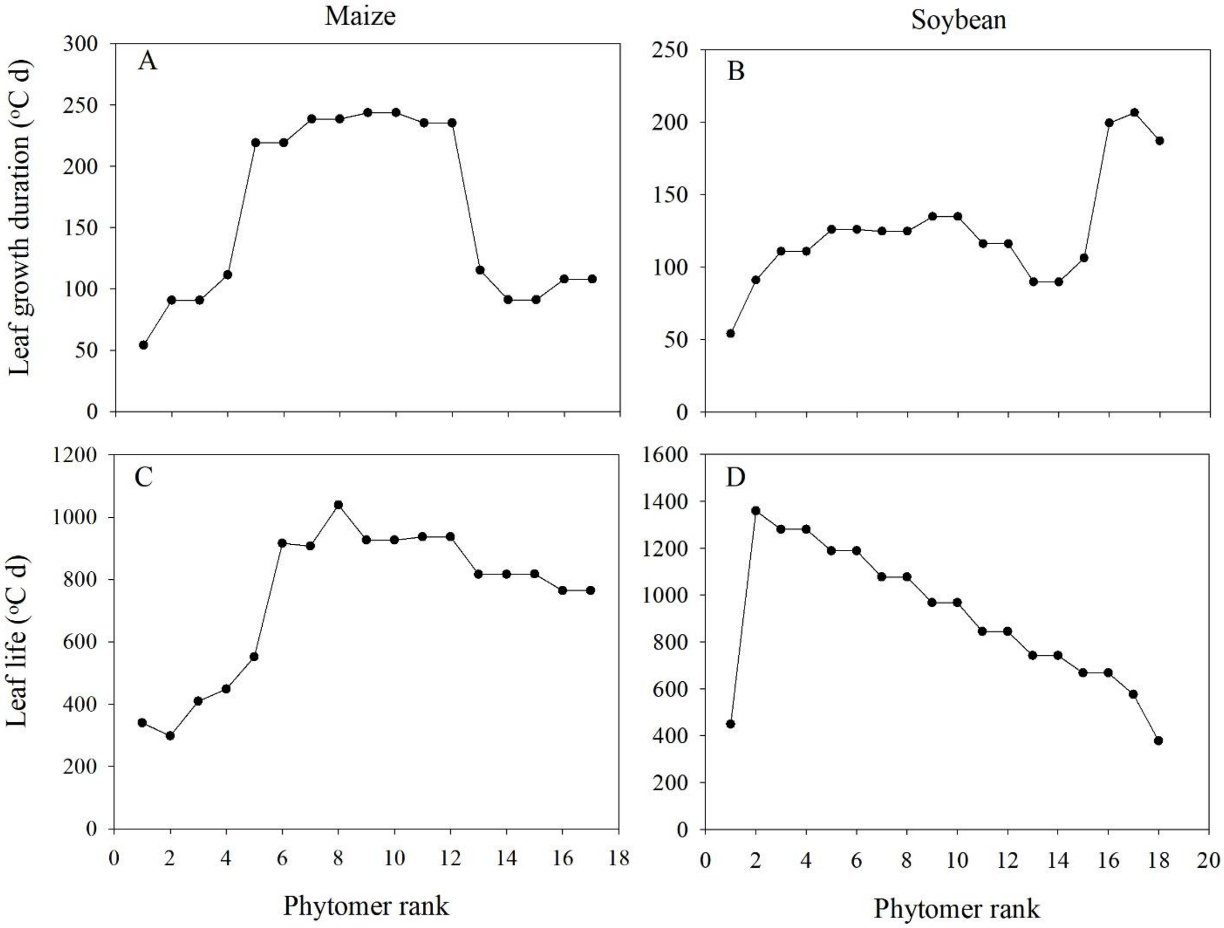


**Supplementary Figure 2.** Mean leaf growth duration ($t_e$) and leaf life across phytomer ranks in monoculture maize (A, C) and monoculture soybean (B, D). Thermal time between leaf appearance and the fully developed phytomer is equal to $t_e$ and the thermal time between the fully developed phytomer and the senescence of the leaf is equal to the leaf life. The average $t_e$ was an input value and the average leaf life value was divided by the average $t_e$ to derive a dimensionless input value.

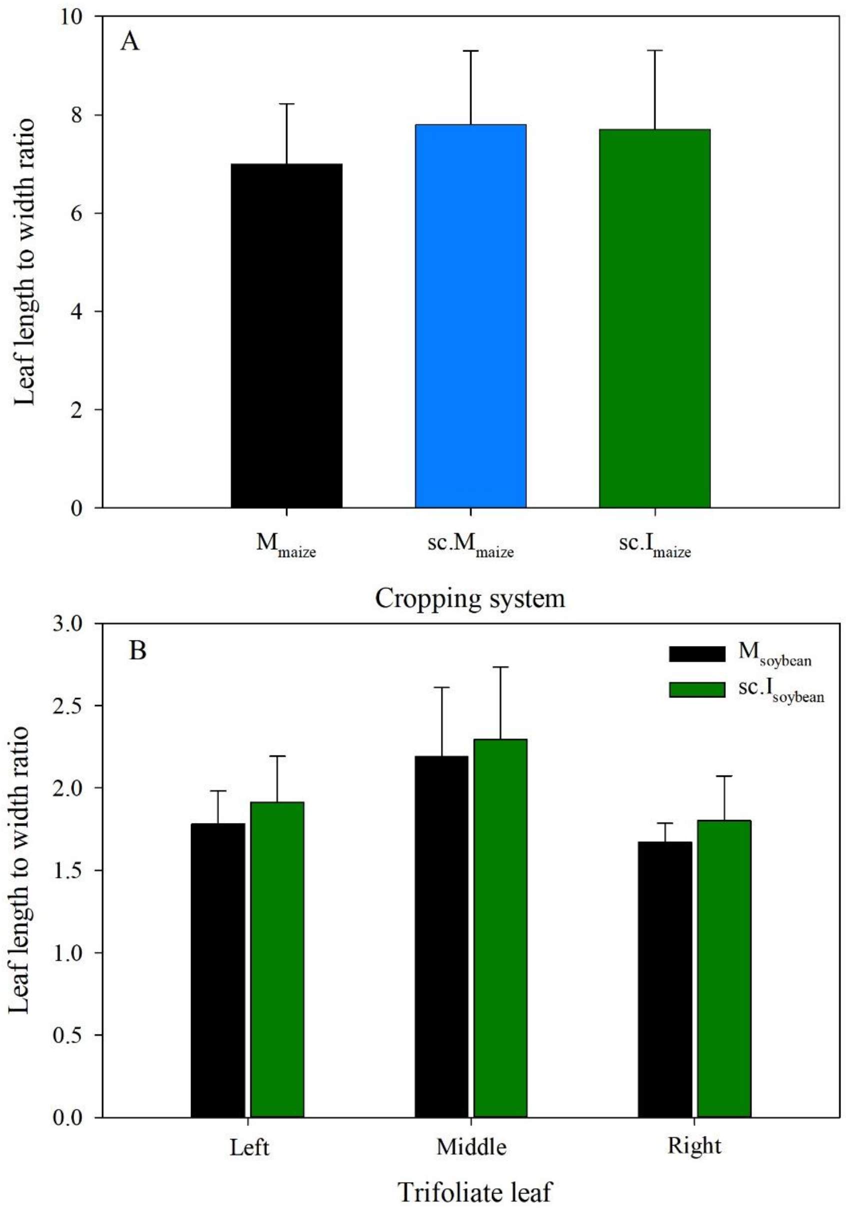


**Supplementary Figure 3.** Mean leaf length to width ratios for maize (A) and soybean (B) under monoculture ($M_{maize}$, $M_{soybean}$), solar corridor monoculture ($sc.M_{maize}$) and solar corridor intercrop ($sc.I_{maize}$, $sc.I_{soybean}$) systems. Values were calculated using a handheld laser leaf area meter (CI-203, CID Bio-Science, USA) across phytomer ranks and the monoculture average was input into the model. Error bars represent the ± SD and n = 4.

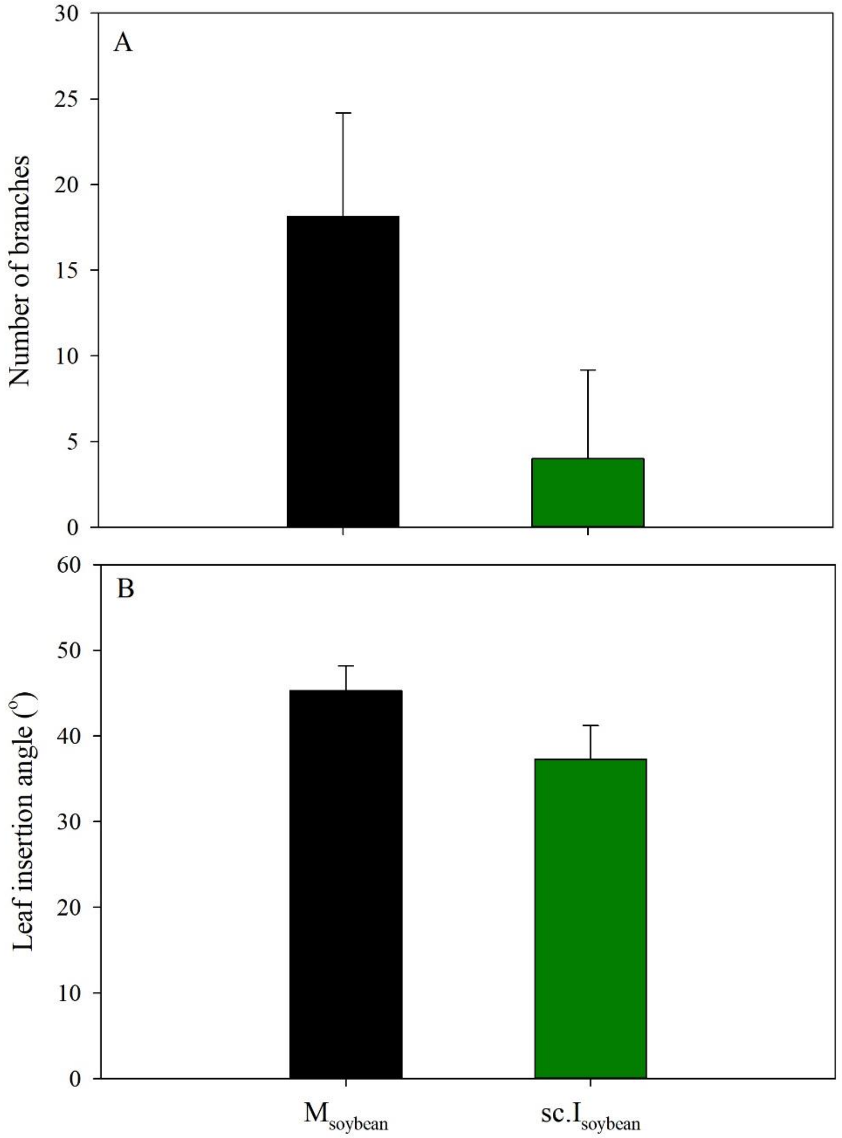


**Supplementary Figure 4.** Mean number of branches (A) and leaf insertion angle (B) for soybean under monoculture ($M_{soybean}$) and solar corridor intercrop ($sc.I_{soybean}$) systems. Leaf insertion values were calculated using ImageJ (https://imagej.net/software/imagej/) on images taken horizontally over plants collected during destructive harvests. Monoculture averages were input into the model. Error bars represent the ± SD and n = 4.

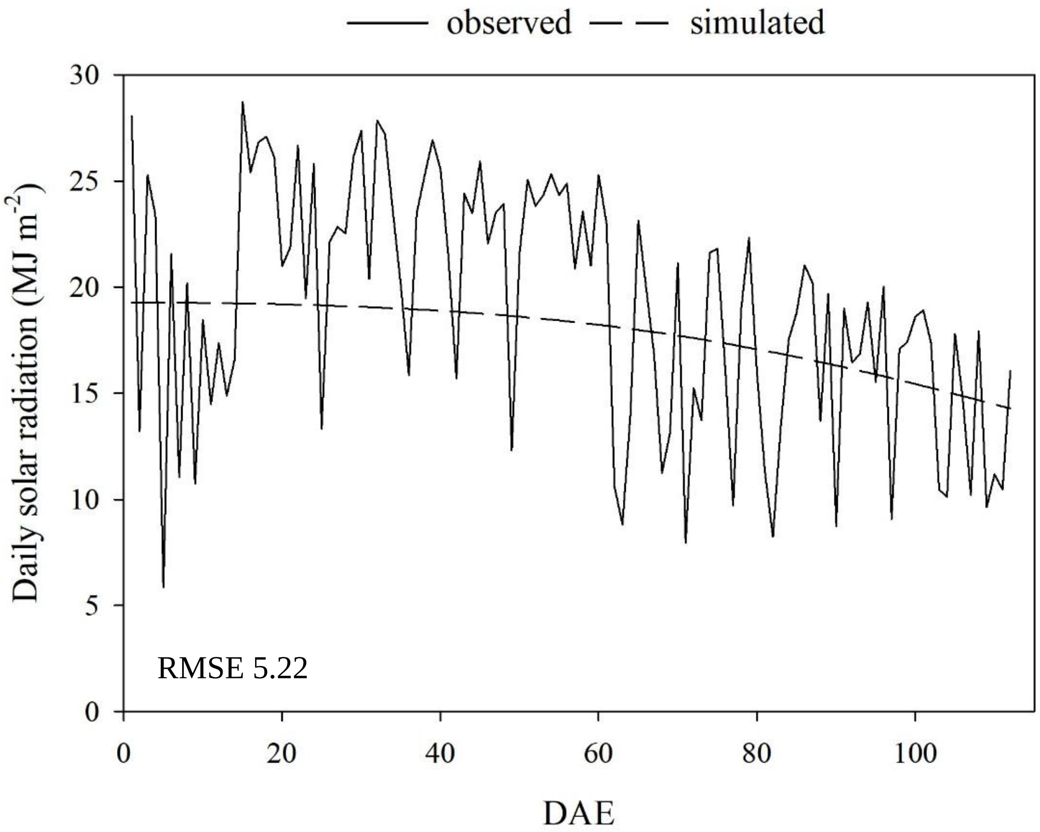


**Supplementary Figure 5.** Mean daily solar radiation observed (solid line) and corresponding simulated values (dotted line) across days after emergence (DAE) during the 2019 growing season. The simulated values were calculated by fitting a sine wave curve and the root mean squared error (RMSE) is reported.

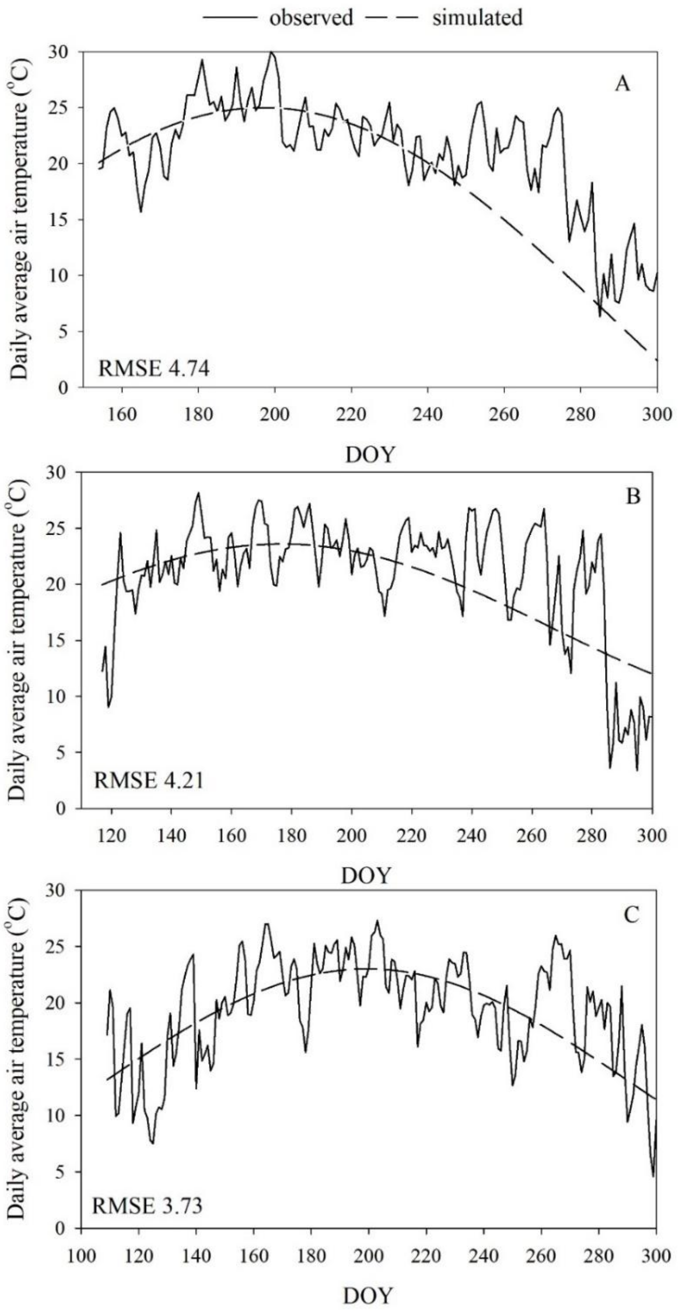


**Supplementary Figure 6.** Mean daily air temperatures at experiment site during 2019 (A), 2018 (B) and 2017 (C) growing seasons (solid line) and the corresponding simulated values (dotted line). The simulated values were calculated by fitting a sine wave curve and the root mean squared error (RMSE) is reported. Only the 2019 values were used for verification and trait analysis simulations. The 2018 and 2017 values were used for validation simulations.

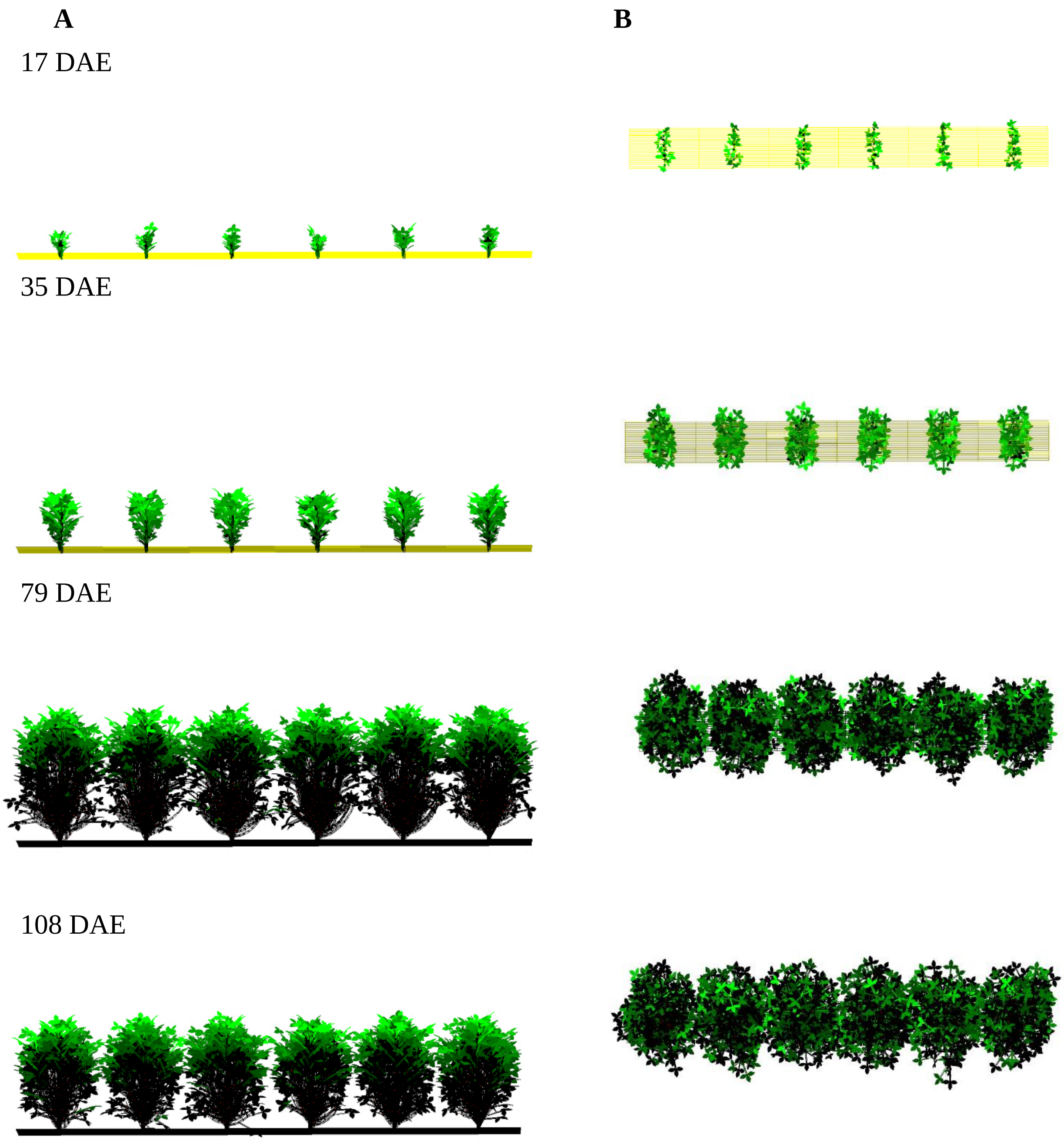


**Supplementary Figure 7.** Comparison of side **A** and bird's-eye view **B** snapshots from soybean monoculture simulations at V3 (17 days after emergence [DAE]), R1 (35 DAE), R6 (79 and 108 DAE) growth stages. All traits were set to the monoculture phenotype. The colour gradients in **B** represent the proportion of light intercepted, where light interception increases from black to green for plant organs and from black to yellow for the soil surface.

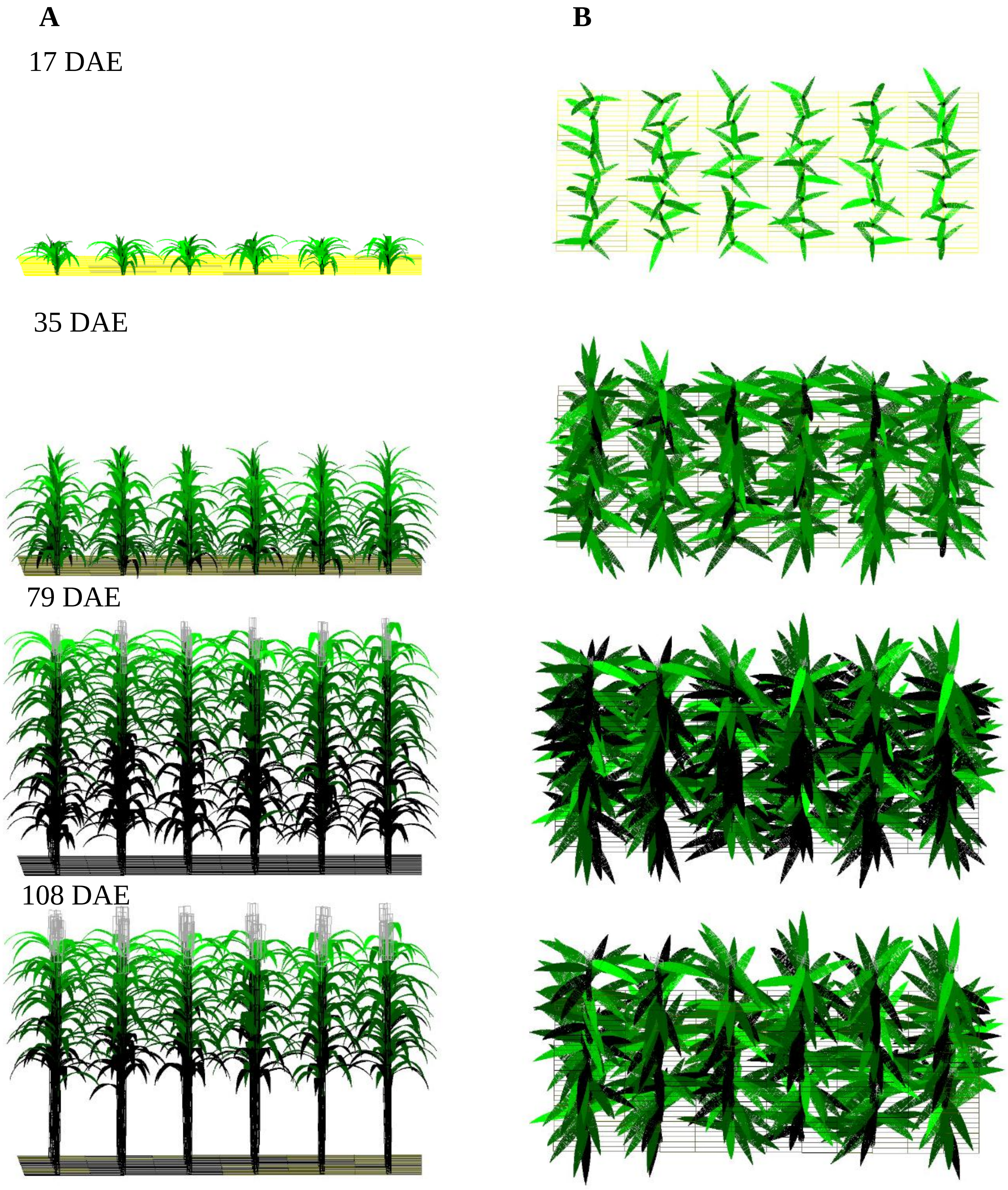


**Supplementary Figure 8.** Comparison of side **A** and bird's-eye view **B** snapshots from maize monoculture simulations at V5 (17 days after emergence [DAE]), V10 (35 DAE), R3 (79 and 108 DAE). All traits were set to the monoculture phenotype. The colour gradients are as in Supplementary Figure 7.

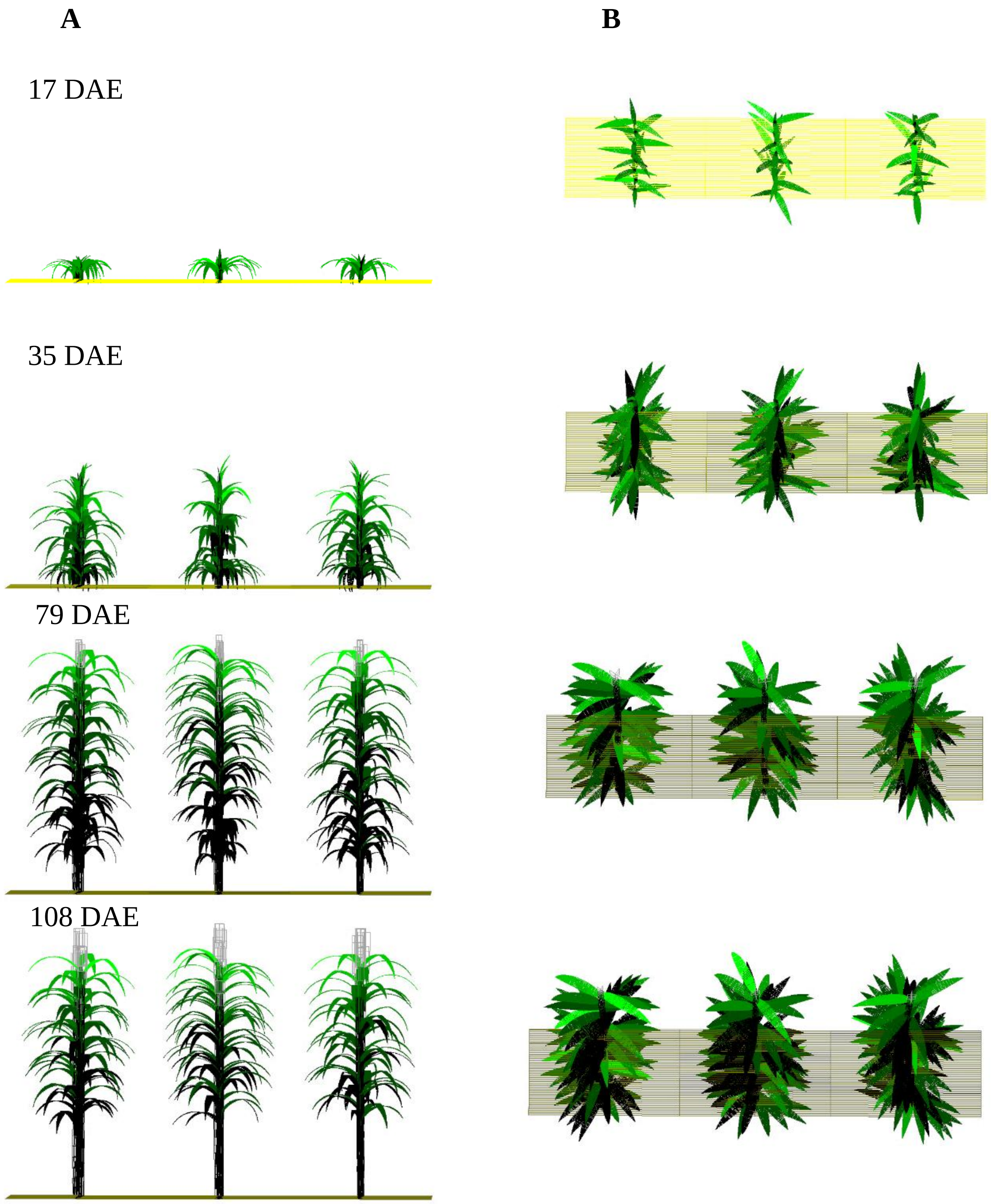


**Supplementary Figure 9.** Comparison of side **A** and bird's-eye view **B** snapshots from maize solar corridor monoculture simulations. All traits were set to the monoculture phenotype. Growth stages are as in Supplementary Figure and the colour gradients are as in Supplementary Figure 7.

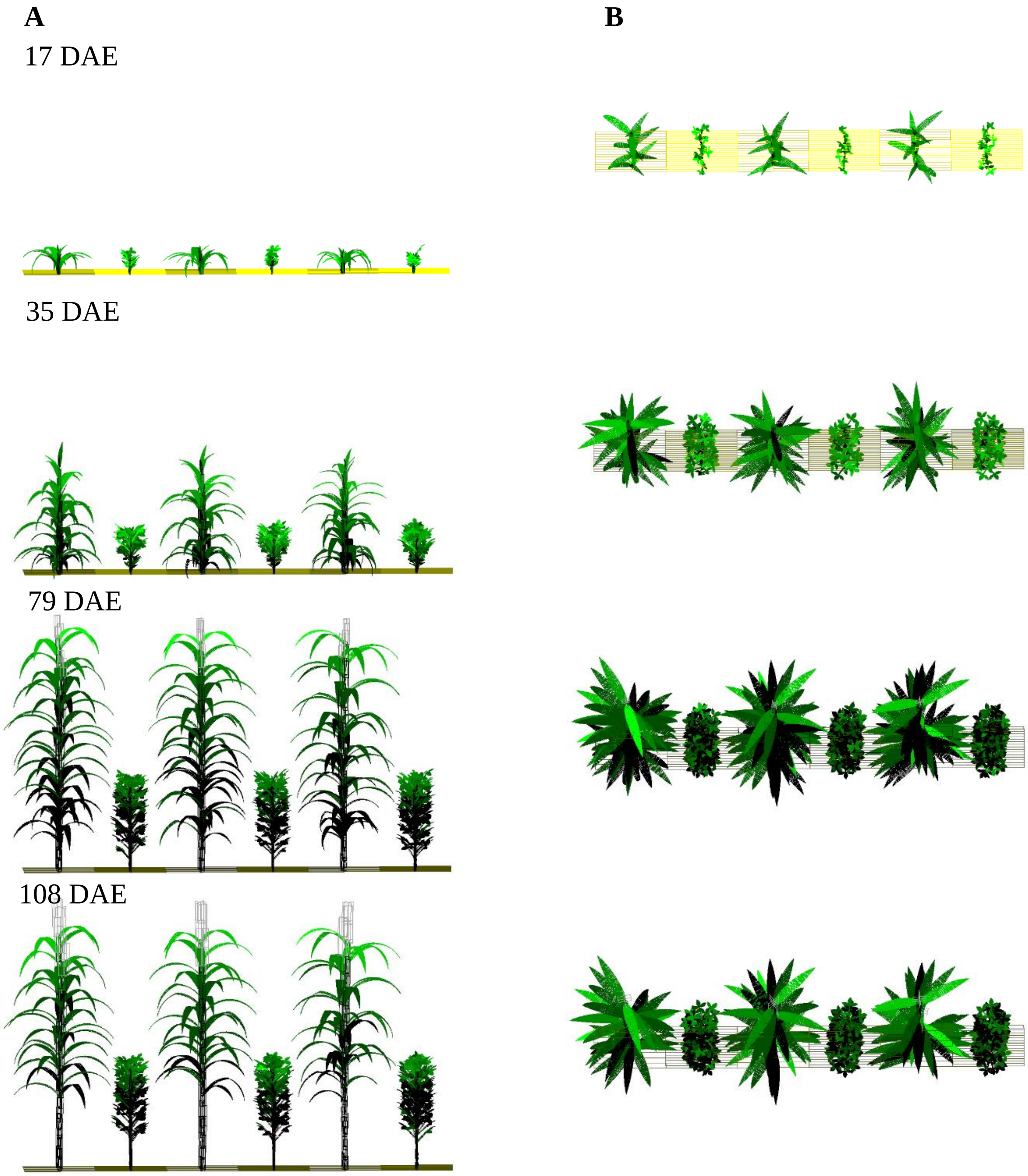


**Supplementary Figure 10.** Comparison of side **A** and bird's-eye view **B** snapshots from solar corridor intercrop simulations at equal growth stages as in Supplementary Figure 8 for maize and Supplementary Figure 7 for soybean. All traits were set to their intercrop phenotype. The colour gradients are as in Supplementary Figure 7.
